# A synthetic gene circuit for imaging-free detection of dynamic cell signaling

**DOI:** 10.1101/2021.01.06.425615

**Authors:** Pavithran T. Ravindran, Sarah McFann, Jared E. Toettcher

## Abstract

Cells employ intracellular signaling pathways to sense and respond to changes in their external environment. In recent years, live-cell biosensors have revealed complex pulsatile dynamics in many pathways, but studies of these signaling dynamics are limited by the necessity of live-cell imaging at high spatiotemporal resolution^1^. Here, we describe an approach to infer pulsatile signaling dynamics from just a single measurement in fixed cells using a pulse-detecting gene circuit. We computationally screened for circuit with pulse detecting capability, revealing an incoherent feedforward topology that robustly performs this computation. We then implemented the motif experimentally for the Erk signaling pathway using a single engineered transcription factor and fluorescent protein reporter. Our ‘recorder of Erk activity dynamics’ (READer) responds sensitively to both spontaneous and stimulus-driven Erk pulses. READer circuits thus open the door to permanently labeling transient, dynamic cell populations to elucidate the mechanistic underpinnings and biological consequences of signaling dynamics.

## Main Text

Many cell signaling pathways exhibit pulses, oscillations or even traveling waves of pathway activity. Examples include the signaling pulses observed from the tumor suppressor p53, the mitogen associated protein kinase (MAPK) Erk, and the immune signaling transcription factor NF-κB^1–5^. Pulses of Erk activity have been observed *in vivo* in the early mouse embryo^6,7^ and in tumors^8^, and self-organize into propagating waves from sites of epithelial injury in both mouse^9^ and zebrafish^10^. The breadth of biological systems exhibiting signaling dynamics suggests that they may play important functional roles. Yet in nearly every context, dynamics are studied exclusively using time-lapse microscopy in single living cells. This granularity of measurement is crucial: to determine whether a cell has pulsed, one must perform at least three measurements to observe a succession of low, high, and low signaling states. However, live imaging can be a severe constraint, limiting the throughput of chemical and genetic screens and restricting *in vivo* studies to tissues that are compatible with single-cell imaging. We thus set out to explore whether we might be able to construct simple synthetic gene circuits to label pulsing cells without live imaging (**Figure 1A**).

**Figure 1.**
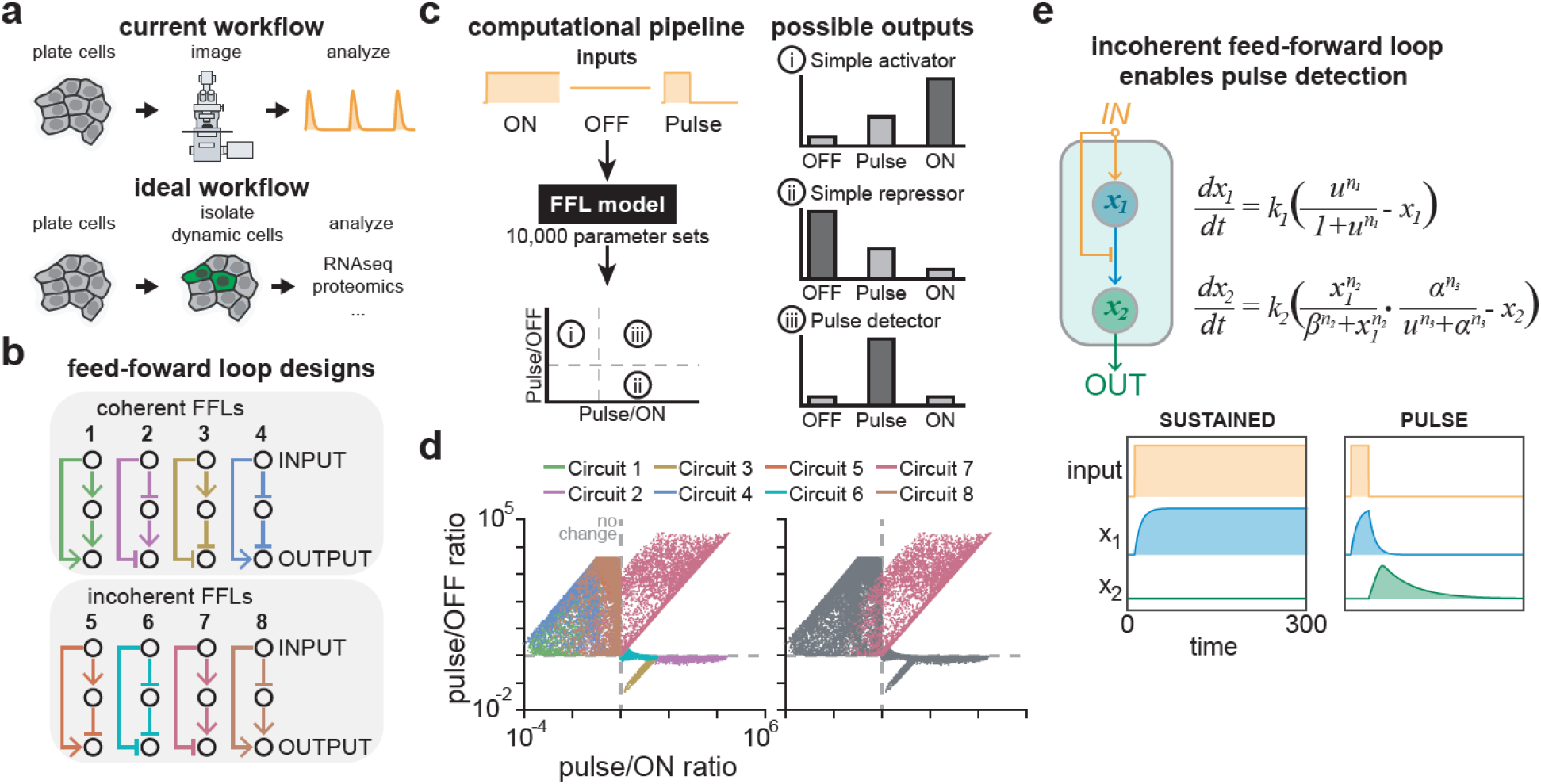
A computational screen for pulse-detecting gene circuits. (**a**) Workflows for studying signaling dynamics. Typically, cells are imaged over time to identify pulses of signaling activity. Ideally, dynamically pulsing cells could be identified using a single fluorescence measurement and then isolated for downstream analysis. (**b**) Coherent and incoherent feed-forward network topologies screened for pulse detection. (**c**) Computational screen workflow: constant ON, constant OFF and pulsed inputs were applied to networks shown in **b** at each of 10,000 random parameter sets. Circuits exhibiting pulse detection would lie in quadrant 3, with pulsed responses greater than constant ON and constant OFF. (**d**) Plot from computational screen with all 8 circuits colored (left), or with only Circuit 7 colored (right). (**e**) Representative time course of Circuit 7 simulated with either constant ON (left) or a pulsed input (right) along with equations describing this circuit for input *u,* intermediate node *x_1_* and output node *x_2_* as described in the inset equation.

Our first goal was thus to identify circuit topologies that might serve as pulse detectors, selectively responding to dynamics while filtering out and ignoring constant high or low signaling states. We focused our attention on feedforward loops (FFLs), a class of network topologies that repeatedly arise in generating or processing dynamic information^11–15^. FFLs are either coherent or incoherent based on whether the two paths connecting input and output have the same or different signs (**Figure 1B**). We devised a simple, modular 2-equation model to represent all 8 FFLs with AND logic at the output node^12^ (**Figure S1, Supplementary Information**), and in each case simulated 10,000 random parameter sets with 3 input conditions: sustained on, sustained off and a pulse of activation.

We assessed the performance of each circuit by calculating integrated output over time in response to each input. We then plotted the ratio of the pulse-induced response to both the constant-on and constant-off cases (**Figure 1C**). By definition, a pulse detector circuit should show stronger induction in response to a pulse than either constant stimulus, leading to high values of both ratios and enrichment in the upper-right quadrant of such a plot, whereas simple activators (circuits that induce gene expression in proportion to the quantity of input signal) would appear in the lower-right quadrant and simple repressors would appear in the upper-left quadrant (**Figure 1C**). Analysis of all 8 FFL topologies revealed that only a single topology, Circuit 7, was capable of performing pulse detection (**Figure 1D**, left). Pulse detection also appeared to be a robust feature of the Circuit 7 FFL, with 96% of simulations showing a stronger response to pulsed stimuli than either high or low constant inputs (**Figure 1D**, right). We also tested all 8 FFL topologies with OR logic at the output node (**Figure S2**). While none exhibited pulse-specific activation, one OR-FFL circuit did exhibit pulse-specific repression and can be understood as the logical inverse of our pulse-detecting Circuit 7 FFL (see **Supplementary Information**; **Figure S2B**).

Examining simulation trajectories provided further insight into the operation of the Circuit 7 FFL (**Figure 1E, Figure S3**). Application of a stimulus (“input”) rapidly results in production of an intermediate node *(x_1_),* but also blocks the ability for *x_1_* to activate an output node *(x_2_).* Only upon removal of the stimulus is repression relieved, enabling *x_1_* to trigger output. Constant-on inputs are unable to trigger a response because input permanently blocks output, whereas constant-off inputs fail because the essential activator *x_1_* is not produced. Overall, our simulations reveal an intuitive and logical relationship between the Circuit 7 FFL topology and pulse detection, demonstrating that pulse detection can arise quite generally out of this particular FFL architecture.

We next set out to implement our pulse detector circuit in the context of a dynamic signaling pathway in mammalian cells, the Erk pathway. Our implementation centered around a single synthetic transcription factor that is regulated by Erk in two opposing ways (**Figure 2A**). For the forward activation path (e.g., Erk input activating *x_1_* which then activates *x_2_*) we envisioned a two-step transcriptional cascade: an Erk-responsive promoter to drive expression of a synthetic Gal4-VP64 transcription factor, which then induces GFP expression from a Gal4-responsive UAS promoter. To match the Circuit 7 FFL topology, our synthetic transcription factor must also be rapidly and reversibly inhibited by Erk (so that the Erk input also directly inhibits *x_2_* production). We realized that fusion with an Erk “kinase translocation reporter” (ErkKTR) would be ideal for implementing this stimulus-dependent inhibition of the engineered transcription factor^16^. Because the ErkKTR is exported from the nucleus in response to Erk activity, an ErkKTR-transcription factor fusion protein would be precluded from encountering DNA and expressing a target gene as long as the pathway remained active.

**Figure 2.**
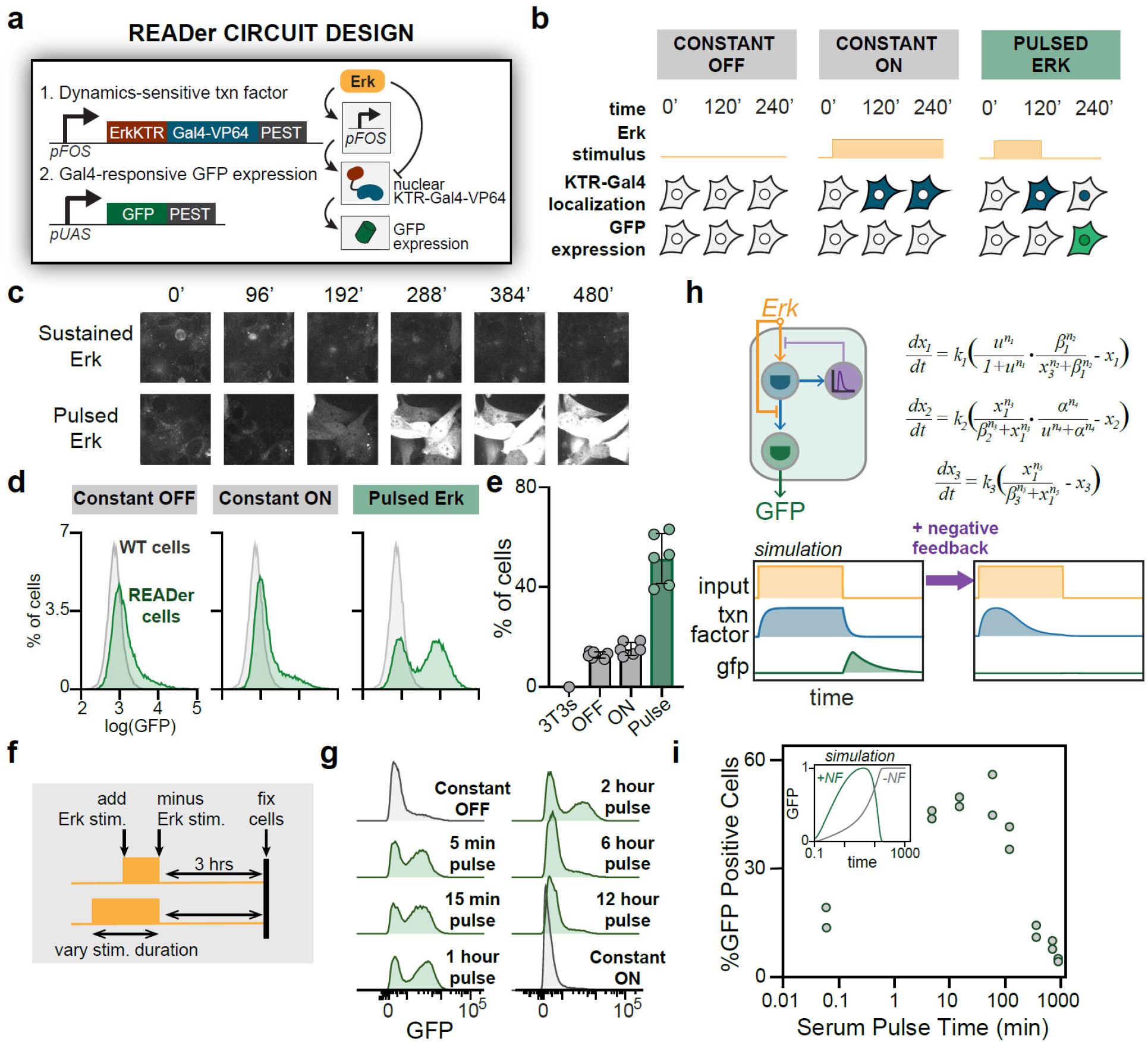
Experimental implementation of pulse detection for the Erk signaling pathway. (**a**) Schematic overview of Recorder of Erk Activity Dynamics (READer). An Erk-responsive promoter drives expression of a KTR-Gal4-VP64 (KGV) fusion protein, which in turn triggers expression of a GFP reporter. (**b**) Illustration of READer circuit logic: (1) under constant OFF stimuli, KGV and GFP levels remain low; (2) under constant ON stimuli, KGV is expressed but exported from the nucleus, preventing GFP production; (3) under pulsed stimuli, KGV is first expressed and then imported into the nucleus, leading to high GFP expression. (**c**) Images of representative fields of NIH3T3 READer cells exposed to constant serum or a 1 h serum pulse. (**d**) Flow cytometry distribution of GFP levels in cells expressing READer (green) incubated in growth factor free media (constant OFF), 10% serum (constant ON) or a one-hour pulse of 10% serum; wild-type NIH3T3s are shown in gray. (**e**) Quantification of flow cytometry data shows % of GFP-high cells in all three conditions. (**f-g**) Mapping how pulse duration affects READer circuit output. Serum inputs of varying duration were applied to cells, which were fixed 3 h after the end of the pulse (schematic in **f**) and analyzed by flow cytometry for GFP induction (data in **g**). (**h**) A extended mathematical model of the READer circuit incorporating previously measured negative feedback on Erk target gene induction. An input *u* (gold) stimulates intermediate node *x_1_* (blue), which produces a negative regulator *x_3_* (purple) that inhibits the production of *x_1_*. (**i**) Quantification of flow cytometry data in **g** reveals that pulses between 5-120 min result in potent GFP accumulation. Inset shows simulated results from the model from **h**, with (green) or without (grey) negative feedback.

To realize this design experimentally we expressed a KTR-Gal4-VP64 synthetic transcription factor (abbreviated throughout as KGV) downstream of the Erk-responsive FOS promoter (P_FOS_). We then used a standard reporter construct, the Gal4-responsive UAS promoter driving destabilized GFP, to record the circuit’s output. Only in response to a pulse of Erk would KGV be first expressed and then shuttled into the nucleus, resulting in GFP production (**Figure 2B**). We termed our circuit – comprising a dynamics-sensitive transcription factor and reporter gene – a Recorder of Erk Activity Dynamics, or READer. We transduced NIH3T3 cells with a lentiviral PUAS-dGFP reporter and transfected them with a PiggyBAC transposase-integrable P_FOS_-KGV Erk-responsive transcription factor, based on our prior data showing that the PiggyBAC system can generate strongly Erk-responsive gene expression^17^, and sorted clonal cell lines harboring both components (**Figure S4**; see **Methods**).

Just as in our simulations, cells expressing the READer circuit were able to discriminate between pulsed and constant signaling inputs. We cultured cells overnight in media lacking growth factors (GF-free media), and then monitored GFP induction by time-lapse microscopy after addition of 10% serum (constant-on), or a 1 h pulse of serum followed by a return to GF-free media (pulsed) (**Figure 2C; Figure S5; Movie S1**). Performing confocal imaging for GFP induction in each case revealed that a pulse of serum led to strong GFP induction within 4 hours, whereas constant-on and constant-off stimuli each led to minimal GFP accumulation.

We reasoned that a pulse detection circuit should also enable inference of prior signaling dynamics from a single measurement in fixed cells. We again exposed NIH3T3 READer cells to constant-off, constant-on and pulsed serum inputs, fixed cells 4 h after the start of stimulation and performed flow cytometry for GFP levels (**Figure 2D**). We found that constant-on and constant-off conditions failed to induce GFP in most READer cells, with a small tail of GFP-high cells that will be discussed in detail below. In contrast, a pulse of serum induced strong GFP induction within 3-6 hours in approximately 50% of cells, while the remainder of the population remained un-induced (**Figure 2E**; **Figure S6**). Subsequent experiments revealed that this bimodal response arose because only a subset of cells could transmit Erk activity to downstream gene expression. Cells sorted from only the GFP-high or GFP-low populations generated the same bimodal response upon a second stimulus challenge, indicating a non-genetic source of response variability (**Figure S7**). Furthermore, the fraction of signaling-responsive cells could be increased by pre-treatment with 10 ng/mL anisomycin (**Figure S8**), a treatment that we previously observed to increase Erk-dependent transcription of endogenous immediate-early genes^18,19^. We also tested whether Erk-triggered target gene induction depended on cell cycle phase, but found that GFP-high and GFP-low cells each exhibited similar DNA content distributions, arguing against cell cycle control over IEG induction (**Figure S9**). Together, our data demonstrates that the READer system labels cells with pulsatile Erk activity and a permissive transcriptional state for immediate-early gene induction.

The scalability of flow cytometry enabled us to rapidly scan additional stimulus conditions to test how the READer circuit filtered a broad range of dynamic inputs. We first tested how selective the circuit was to changes in the pulse duration. Endogenous Erk pulses are typically observed to be less than 1 h in length, with sustained responses lasting for multiple hours^8,20–22^. We applied pulses of different durations ranging from 5 min to 12 h, then incubated cells for an additional 3 h prior to fixation to allow GFP to accumulate (**Figure 2F**). Although pulses from 5 min to 2 h resulted in similar profiles of GFP expression, longer pulses were filtered and ignored by the circuit (**Figure 2G**). We also tested for GFP induction in response to a broad range of dynamic Erk inputs delivered using our OptoSOS optogenetic system (see **Methods**)^23,24^. Trains of multiple pulses also led to GFP accumulation, indicating that the READer circuit detects persistent signaling oscillations as well as a single pulse (**Figure S10**). Together, these data reveal that the READer circuit responds broadly to pulsatile Erk stimuli while filtering out constant high or low signaling states.

Our data revealed that the READer circuit ignores very long pulses greater than 2 h in length (**Figure 2G**). While this long pulse rejection was not a prediction from our original “Circuit 7” model, it can be readily understood based on the prior observation that even a sustained Erk stimulus only drives a transient, 30 min pulse of IEG expression, after which subsequent expression is suppressed^25–27^. Thus, after a long input pulse, KGV RNA/protein levels could drop, leaving little protein to return to the nucleus to drive GFP expression (**Figure 2H**). We verified that our transposase-integrated P_FOS_ promoter indeed produced a transient pulse of expression, in agreement with prior data on *fos* expression (**Figure S11**)^25^. Implementing transient P_FOS_-driven expression in our computational model was also sufficient to match the duration-based filtering that we observed experimentally (**Figure 2I**, inset). To further probe this framework, we queried the model for parameters that might tune pulse detection (**Supplementary Information; Figure S12-13**). Our parameter scans indicated that destabilizing the KGV transcription factor could further shift READer sensitivity to shorter-duration pulses, a prediction we confirmed experimentally by incorporating destabilized 3’UTR and PEST sequences on the KGV mRNA and protein^28,29^. Indeed, cells with destabilized KGV variants only induced GFP in response to pulses of 1 h or less (**Figure S14**). Overall, our simulations and experiments converge on an intuitive result: the READer circuit acts as a band-pass filter whose pulse detection characteristics can be further turned by modulating the mRNA/protein stability of the engineered transcription factor.

We have seen that the READer circuit responds selectively to a stimulus pulse; can it also detect spontaneous, naturally occurring Erk pulses? We noticed that a small subpopulation of READer-expressing fibroblasts expressed high levels of GFP even when cultured under constant stimulus conditions (**Figure 2D**), raising the possibility that this sub-population may undergo spontaneous Erk pulses that are then detected by the READer circuit. To directly confirm whether such a pulsatile sub-population exists, we transduced NIH3T3 fibroblasts with a fluorescent ErkKTR-irFP biosensor and imaged them for 48 hours under continuous serum and GF-free conditions. Indeed, we observed that some cells began to pulse spontaneously after approximately 12 h of culture (**Figure S15; Movie S2**). These data would be consistent with the appearance of a GFP-positive population being driven by spontaneous pulses.

To directly compare endogenous Erk pulses to GFP accumulation, we next transduced our READer clonal cell line with the ErkKTR-mScarlet fluorescent biosensor to monitor both biosensors in the same live cells (**Figure 3A**). We incubated cells in serum-free media overnight and switching to ‘constant-on’ growth media at the start of imaging, based on simulations which indicated that such an input should prevent READer system activation during the constant-on phase but elicit a sharp rise in GFP induction upon the spontaneous switch to pulsatile Erk activity (**Figure 3B**). Indeed, we found that serum stimulation first drove a constant-on Erk state (leading to ErkKTR nuclear export), but after 15 hours Erk activity began pulsing in some cells (**Figure 3C; Movie S3**). The switch to a pulsing state was accompanied by a rapid increase in GFP intensity, a phenomenon that was matched by our computational model when stimulated with the experimentally-observed trajectory of Erk dynamics (**Figure 3C, inset**; see **Figure S16** for additional cells and simulations). Taken together, these data demonstrate that the READer circuit can indeed sense spontaneous, naturally occurring Erk pulses.

**Figure 3.**
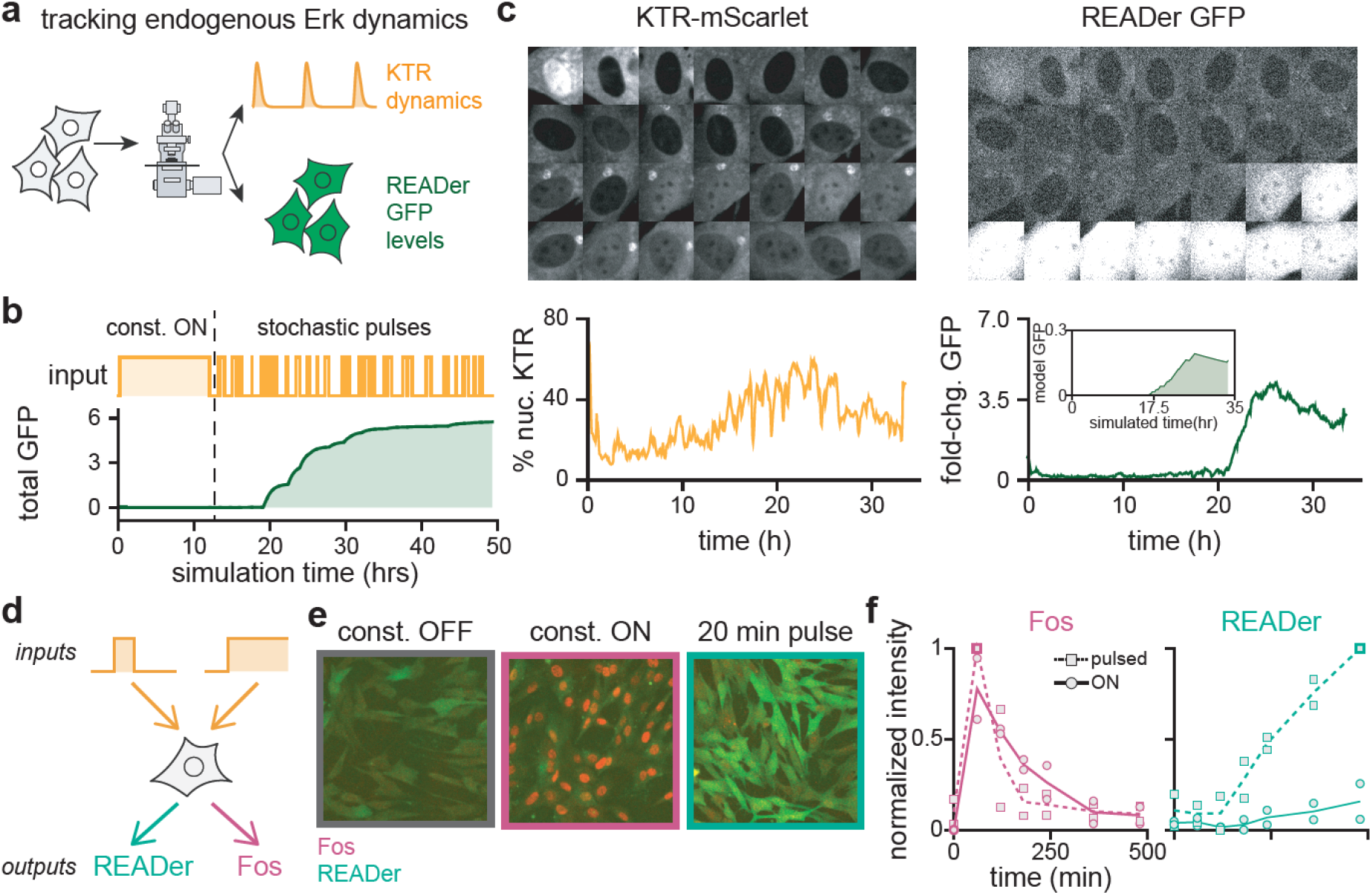
READer detects endogenous Erk pulses and encodes a distinct axis from class Erk target genes. (**a**) Combining direct measurement of endogenous Erk dynamics with the READer system. NIH3T3 READer cells transduced with KTR-mScarlet can be used to simultaneously visualize Erk activity dynamics and READer GFP output in single cells. (**b**) Simulated cellular response during the switch from constant ON to stochastic pulses of Erk activity. The simulated GFP response from the mathematical model of Figure 2I is shown (green). (**c**) Images and quantification from a representative READer cell expressing KTR-mScarlet stimulated with serum at time 0 and imaged for 34 hours. Confocal images of nuclear KTR-mScarlet and GFP levels are shown (top), with quantification of nuclear KTR and fold-change in GFP intensity (bottom). Inset shows simulated GFP response when the same pulsatile KTR-mScarlet trace is used as a model input (see Methods). (**d**) Comparing simultaneous imaging of READer and the canonical Erk target gene Fos in fixed cells to determine whether READer provides orthogonal information. (**e**) Representative images of NIH3T3 READer cells that were incubated in growth factor free media (constant OFF), 10% serum (constant ON) or a 20 min pulse of serum (pulsed), fixed, and imaged for GFP (green) and Fos (red). (**f**) Quantification of immunofluorescence intensity from fixed cells as in **e** for Fos (left) and READer GFP (right). Solid line shows sustained serum stimulation; dashed line shows the 20 min serum pulse. Error bars represent the standard error of the mean for biological duplicates.

As a final test of the READer biosensor, we set out to compare its response to staining for classic Erk target genes. To our knowledge, no endogenous Erk target genes have been identified that specifically sense pulsatile stimuli, but the Fos immediate-early gene product has long been used as a marker to identify cells exhibiting high levels of Erk pathway activity^30^ (**Figure 3D**). Fos staining has been particularly useful in neuroscience, as it labels cells that have recently experienced high levels of neuronal activity^31,32^. To directly compare Fos and READer activation in response to different dynamic stimuli, we incubated cells overnight in GF-free media, and switched them to either sustained growth media or a 20 min pulse of growth media. We then fixed cells at various time points post-stimulus and monitored Fos protein by immunofluorescence and READer-induced GFP fluorescence in the same cells (**Figure 3E-F**; see **Figure S17** for full joint READer/Fos distributions). We observed rapid, strong induction of Fos at early time points regardless of stimulus duration, demonstrating that simply measuring Fos cannot be used to discriminate pulsatile from sustained signaling. In contrast, the READer circuit only triggered GFP expression in response to a pulse but not constant stimulation. These results confirm that the dynamic information using READer cannot be obtained by staining for classic Erk target genes like Fos.

Here we report the discovery and characterization of a simple gene network that can selectively and robustly differentiate between pulsatile and constant signaling states. Our network is based on an incoherent feedforward loop with slow activation and fast repression. Incoherent feedforward loops have been studied extensively for their dynamic filtering capabilities, including pulse generation and temporal ordering^11,12,15^; our work adds highly-selective pulse detection to this list of capabilities. We also report a simple, flexible implementation of this network architecture for mammalian signaling, centered on the pathway-regulated expression of a transcription factor that is fused to a kinase translocation reporter. Although we have focused on Erk signaling in this work, we believe this report provides the roadmap to the development of a suite of new reporters that capture the previous dynamic history for many dynamic signaling pathways (i.e. p53, Wnt, NFkB, etc.)^2,4,5,33,34^. Kinase translocation reporters are available for a growing number of proteins and pathways^35,36^ and other forms of fast negative regulation (e.g. signal-induced protein degradation) would be expected to work with similar efficacy. Biosensors like the READer circuit could be transformative for mapping signaling dynamics *in vivo*, for large-scale genetic screens to identify the biochemical networks that generate pulses, and for tracing the lineages and eventual fates of pulsing cells. Such circuits may shed new light on the roles played by signaling dynamics in diverse contexts from disease to development.

## Author contributions

Conceptualization: P.T.R., and J.E.T.; Methodology: P.T.R. and J.E.T.; Experimental Investigation: P.T.R. Mathematical Modeling: P.T.R., S.M. and J.E.T. Data analysis: P.T.R., and J.E.T.; Writing – Original Draft: P.T.R., S.M. and J.E.T.; Writing – Review and Editing, all authors; Funding Acquisition: J.E.T.; Resources: J.E.T.; Supervision: J.E.T.

## Acknowledgements

We thank all members of the Toettcher lab for helpful comments. We also thank Katherine Rittenbach and Dr. Christina DeCoste of the Princeton Molecular Biology Flow Cytometry Resource Center for cell sorting. This work was supported by NIH grant DP2EB024247 (to J.E.T.), the Hertz Foundation Fellowship and NSF GRFP Fellowship (to S.M.) and the Lidow Independent Work 2019 Research Grant (to P.T.R.).

