## Supplementary Information for "A synthetic gene circuit for imaging-free detection of dynamic cell signaling"

**This PDF file includes:**

| | |
|---|---|
| Online Methods | Pages 2-11 |
| Supplementary Text | Pages 11-29 |
| Supplementary Sequences | Pages 29-30 |
| Supplementary Figures 1-17 | Pages 31-48 |
| Supplementary Movie 1-3 Legends | Page 49 |
| Supplementary References | Page 49-50 |

### Online Methods

#### Plasmid Construction

We cloned all of our constructs/synthetic gene circuits into the pHR lentiviral expression plasmid[1] or into Piggybac synthetic immediate early gene (synIEG) plasmid[2,3]. All linear DNA fragments were prepared by PCR, using GXL polymerase for fragments > 7500 bp and HiFi polymerase (Takara Bio) for shorter fragments. After PCR, Dpn1 digestion was used to remove template DNA. PCR products were cut from agarose gels and purified using the Nucleospin gel purification kit (Takara Bio). Final plasmids were constructed using Infusion assembly and amplified in Stellar chemically competent *E. coli* (Takara Bio). DNA was extracted by miniprep (Qiagen). All plasmid verification was done by restriction enzyme digestion and Sanger sequencing (Genewiz).

To create the KTR-Gal4-VP64 synthetic transcription factor plasmid, we took the original synIEG plasmid (pFos – destabilized GFP – Tubulin 3'UTR – pCMV – BFP) and replaced the GFP with KTR-iRFP (Addgene #111510)[4], keeping the PEST destabilizing element at the end of the coding region. In a second round of cloning, the iRFP was replaced with a Gal4-VP64 fusion protein amplified from pHR_pGK_LaG17_synNotch_Gal4VP64 (Addgene number: 79127, a gift from the Lim lab)[5]. For the reporter gene, we started pHR_Gal4UAS_IRES_mC_pGK_tBFP (Addgene #79123)[5]. First, the tBFP was replaced with iRFP (Addgene# 111510)[4]. In a second round of cloning, we amplified destabilized GFP from the original synIEG plasmid and cloned it in place of the IRES-mCherry element.

#### READer cell line generation

*Lentivirus production*

HEK 293Ts were plated in a 6 well plate at ~40% confluency at least 12 hours before transfection. The cells were then co-transfected with 1.5 ug of the pHR reporter vector along with 1.33 ug and 0.17 ug of CMV and pMD packaging plasmids, respectively, using Fugene HD (Promega). Virus was collected after 48 hours post-transfection and filtered through a 0.45 μm filter. To the ~2 mL of viral media, 2 μL of polybrene and 40 μL of 1 M HEPES were added. Cells to be infected were plated at 40% confluency in a 6 well plate at least 12 hours before infection and then 200-500 μL of viral media was added the cells. 24 hours post transduction virus containing media was replaced with fresh media.

*PiggyBac Integration*

NIH 3T3s that were infected with the reporter construct and were to be integrated with dynamic TF PiggyBAC plasmid were plated 24 hours prior to transfection. 2.08ug of the plasmid to be integrated along with 0.41 μg of the PiggyBAC helper plasmid were co-transfected into the cells using Liptofectamine LTX with Plus reagent. Cells were then selected using FACS for dual BFP/iRFP expression after 3 days post-transfection. Cells underwent single cell sorting to isolate clonal cell lines.

*Variants on cell line*

To perform optogenetic experiments, pHR OptoSOS-tgRFP was introduced into clonal cell line generated above. To do so, lentivirus was generated from pHR SFFVp tgRFP-SSPB-SOScat-P2A-ILID-CAAX as described above. READer clonal cells were plated at ~40% confluency in 6-well plates and infected with virus. To perform simultaneous visualization of

Erk pulsatile activity and READer output, we lentivirally infected the original READer clonal cell line with pHR SFFVp KTR-mScarlet.

To develop the versions on the READer circuit with different degradation properties, we first generated a clonal cell line containing the OptoSOS construct as well as the reporter plasmid (both introduced lentivirally). This line was then validated to have photoswitchable Erk activation by introducing KTR-iRFP using lentivirus[6] and applying cycles of light. After this, 4 different variants of the READer circuit (combinations of PEST/PESTmut and *tubulin/fos* 3'UTR) were introduced using PiggyBAC integration method described above and cells were subjected to another round of clonal cell line generation.

**Cell line maintenance and preparation for imaging**

All cells (NIH 3T3s and HEK 293Ts) were grown in DMEM plus 10% FBS in Thermo Fischer Nunc Cell Culture Tissue Flasks with filter caps at 37C and 5% $CO_2$. Cells to be imaged were plated into InVitro Scientific's 96 well, black-walled, 0.17mm high performance glass bottom plates. 10 µg / mL of fibronectin diluted in PBS was placed on the wells, washed off and then cells were plated in DMEM with 10% FBS for at least 12 h. 12 h prior to imaging, cells were placed in growth factor free media (DMEM with 0.00476 mg / mL HEPES). 50 µL of mineral oil was pipetted onto the wells right before placing onto the scope to prevent media from evaporating.

**Imaging**

Cells were maintained at 37C with 5% CO2 for the duration of all imaging experiments.

Confocal microscopy was performed on a Nikon Eclipse Ti microscope with a Prior linear motorized stage, a Yokogawa CSU-X1 spinning disk, an Agilent laser line module containing 405, 488, 561 and 650 nm lasers, a 60X oil or 20X air objective and an iXon DU897 EMCCD camera.

**Preparation of fixed cell samples**

Cells to be analyzed via flow cytometry were plated at ~60% confluency in 6-well plates. 36 hours post-plating and 12 hours before the experiment began, cells were switched into growth-factor free media (DMEM with 0.00476mg/mL HEPES). Serum additions (up to 10%) were then given and removed depending on the stimulus. After stimulation protocol was applied, media on cells was aspirated and cells were washed with PBS. After trypsinizing cells, neutralizing the trypsin with media and spinning cells down, the supernatant was removed and the cell pellet was resuspended in a 2% PFA solution (50:50 mixture of BD CytoFix solution and PBS). After leaving the samples at 4 °C for 20 minutes, cells were spun down and then resuspended in cold PBS. Samples were run on LSRII Flow cytometer between 24 and 48 hours after fixation. Gating strategy is described in the supplemental text.

**Pulse experiments**

*Simple, single pulse experiment*

READer clonal cells were plated into 6 well plates 24 hours before the start of stimulation and placed into growth-factor free media 12 hours prior to stimulation. For the constant off condition, cells were given 200 µL of growth-factor free media and both the constant on and pulsed conditions were given 200 µL of FBS (to a final concentration of 10% v/v). After 1 hour, the media on the cells of the pulsed and constant off condition was removed, and fresh growth-factor free media was added while the media for the cells in the constant on condition was switched for regular growth media (containing growth factors). After 3 hours, the cells underwent the fixation protocol described above.

*Bandpass pulse experiment – serum*

READer clonal cells were plated into 6 well plates 24 hours before the start of stimulation and placed into growth-factor free media 12 hours prior to stimulation. After this 12-hour starvation period, the constant on condition was given serum to a final concentration of 10% v/v, and the longest duration of pulse (typically 12 hours) was given serum to the same final concentration. Then, working backwards from longest pulse duration to shortest, serum was added to the appropriate well. After the shortest pulse duration, all wells that were receiving pulsed inputs, as well as the constant off condition, were placed in growth-factor free media while the constant ON condition was placed in full growth media for 3 hours. After this wait time, cells underwent the fixation protocol described above.

For the optogenetic bandpass experiments described in Figure S10, READer clonal cells transduced with the OptoSOS system were plated in individual 35mm dishes 24 hours before the start of the experiments and, upon being placed into growth factor free media 12 hours prior to stimulation, were placed in a box covered in foil to protect the cells from the light. As described above, in order from longest pulse duration to shortest, the individual dishes of cells were placed in a light box stimulated with blue light. After all of the cells were placed in the box for the appropriate amount of time, the light plate was turned off for 3 hours. After this wait time, cells underwent the fixation protocol described above.

*Off-time analysis*

READer clonal cells were plated into 6-well plates 24 hours prior to the start of stimulation and placed into growth-factor free media 12 hours prior to stimulation. All cells were then given at 15 minute pulses of serum, and subsequent switch to growth-factor free media, at staggered time points. The cells that were going to have the longest off time were given their pulse first, and then went down to the shortest off times. From there, the cells underwent the fixation protocol described above.

*Pulse train experiment*

For pulse train experiments involving optogenetic stimuli, READer cells expressing the OptoSOS system were plated into 6 well plates 36 hours prior to experimentation. 12 hours prior to the beginning of the experiment, cells were placed in growth-factor free media and wrapped in foil to prevent light exposure. Light was delivered using custom-printed circuit boards of blue 450 nm light-emitting diodes (LEDs). During light stimulation, cells were maintained in an incubator at 37 °C in separate foil-wrapped boxes covered with separate blue LED boards delivering different patterns of light inputs. Each LED board was connected to a separate constant-current LED driver, all of which were controlled using an Arduino MEGA 2560 microcontroller board. The Arduino was programmed with open-source IDE software to deliver different dynamic light input regimes to each circuit board. To minimize phototoxicity, light inputs were delivered in cycles of 20 sec ON and 10 sec OFF which allowed us to minimize light exposure while still delivering a constant stimulus to cells by taking advantage of the slow

(~0.5–1 min) dark decay rate of iLID activation[7]. After 12 hours in the light box, the cells underwent the fixation protocol described above.

***Fos* immunofluorescence staining**

READer clonal cells were plated in wells of a 96-well plate. 24 hours post plating, and 12 hours prior to stimulation began, cells were placed into growth-factor free media. 15 minutes prior to each timepoint, the pulse condition was given a 15-minute pulse of serum by adding 10ul (10% by volume final concentration) of FBS and then placed back into growth-factor free media. At the endpoint of the 15-minute pulse, the corresponding 'sustained-input' well for that timepoint was given 10 μL of FBS (10% by volume final concentration). After all of the timepoints were completed, all cells were fixed by removing the media, adding 75 μL of CytoFix solution and incubating at room temperature for 10 minutes. CytoFix was then removed and an additional 100 μL of PBS was added and then dumped out to further wash out remaining CytoFix. Cells were then permeabilized by adding 100 μL of ice-cold 90% methanol to each well and incubating at -20C for 10 minutes. Methanol was then removed and the cells were washed with 100 μL of PBS. Cells were then incubated in IF buffer (PBS + 10% FBS + 2mM EDTA) at room temperature for 1 hour. After blocking, the cells were then placed in 75 μL of primary antibody (Fos antibody Rabbit mAb – 9F6) at a 1:3000 dilution in IF-T buffer (50 mL of IF buffer + 150 μL Trixon X-100) and incubated overnight at 4 C. After primary incubation, cells were washed with IF-T buffer three times with 5-minute incubations between washes. Cells were then placed in 70 μL of secondary antibody that was diluted 1:500 in IF-T buffer and were incubated at room temperature for 1 hour. After 3 more washes using IF-T buffer with 5-minute incubations, DAPI was added to stain nuclei, and PBS was added for long term storage at 4 °C.

Images were acquired with 24 hours of staining.

**Cell cycle analysis**

READer clonal cells were plated into T75 flask at 40% confluency. After 36 hours, the media was switched out for growth-factor free media for 12 hours. Serum was added for a final concentration of 10% by volume for 1 hour and then switched back to growth-factor free media. Cells were then fixed using the protocol delineated above. After 24 hours, the cells were sorted into two tubes, one for GFP-positive and the other for GFP-negative cells using the FACS Aria. The two populations were then spun down and re-suspended in FACS staining buffer (0.1% Triton X-100 in PBS). After a mild vortex to get single-cell suspension, PI was added to a final concentration of 20 μg/mL and RNAase A was added to a final concentration of 200 μg/mL. After a mild vortex, the cells were incubated at room temperature for 30 minutes and then run on LSRII flow cytometer. FSC Express 7 was then used to perform cell-cycle analysis.

**Microscopy Data Analysis**

*Initial READer cell experiment analysis*

ND2 files from Nikon Elements software were imported into ImageJ. The measure tool was used to quantify mean intensity of the nuclei of cells of interest. These files were saved and then imported into R to do statistical analysis and graphing. The code for this analysis can be found in Supplementary code from Ravindran et al[3].

*Simultaneous KTR and READer analysis*

ND2 files from Nikon Elements software were imported into ImageJ. The measure tool was used to quantify mean intensity of the nuclei and cytoplasmic intensity for cell of interest in the KTR channel (mScarlet). These text files were then saved and imported into MATLAB to capture nuclear intensity for READer channel (GFP) and to produce figures. Specifically this code first reads in the two text files that provide the nuclear and cytoplasmic intensities of the KTR channel as well as provides the details for the position of the nucleus. From here, the percent nuclear KTR is calculated by dividing the nuclear intensity by the sum of the nuclear and cytoplasmic intensity and multiplying by 100. The details of the ellipse over time used to quantify the nuclear intensity is then read out (specifically the x and y coordinates, the width and the height of the ellipse). These are then used to create a mask that is then applied on a tif file version of the GFP images such that all intensity values are NaN except for those within the mask. The average intensity is then used to calculate the average GFP intensity in the nucleus over time. Normalized GFP is calculated by subtracting off the background intensity from all time points and then dividing all intensities by the intensity at the first time point.

To feed the KTR intensities into our mathematical model, we first binarized the traces by taking any value above 48% nuclear KTR to mean an Erk off state (input = 0) and any value above below 48% nuclear KTR to mean an Erk on state (input =10). These traces were then fed into the model and the resulting GFP curve (variable $x_2$) was plotted.

*Fos analysis*

ND2 files from Nikon were imported into ImageJ and saved as '.tif' files. These were then imported into MATLAB and custom code was written to analyze the cells. Essentially, masks were created using DAPI image for each condition and then this mask was applied to the

Fos and READer images. The mean nuclear intensity from each cell was then used for downstream analysis.

**Transcriptional Burst Analysis**

Protocol was adapted from Wilson et al., 2017. Briefly, 7 z-stack slices spanning 4.5 um (0.8um between z-slices) which was centered on the middle of the nucleus. This z-stack was max projected to allow all of the bursts to be visualized on a single plane. Positional information was tracked using the measure tool in Fiji. MATLAB code was used to take in the positional information, fit a 2-dimensional Gaussian to the identified region and finally calculated the integrated area under the fitted Gaussian as the burst intensity. The code for this analysis can be found in Supplementary Code from Ravindran et al[3].

### Supplementary Text

##### 1. Computational screen for pulse detecting feed forward loops

##### 1.1 Goal: identification of synthetic gene circuit to respond selectively to pulsatile signaling

Our objective in the report was to find a network architecture that could implement pulse detection in the hopes that we could use this architecture to make a synthetic gene to respond to pulsatile signaling events. With the advent of live-cell biosensors, it has become clear that many signaling pathways do not follow the textbook dogma of activating to a high constant manner. Instead, signaling pathways in mammalian cells display ornate time-varying patterns of activation. More recent advances in optogenetics and microfluidics have shown that different cellular fates rely upon these time-varying inputs to drive specific cell fates. Although connections between dynamic signaling pathways and their consequences are beginning to be revealed, there are still major unanswered questions about where such dynamics may exist *in vivo* and what genetic networks enable such pulsatile activity. As one needs high resolution, long-term imaging to capture dynamic signaling activity, live-cell biosensors are still limited in addressing such questions as they are relatively low-throughput and many endogenous contexts are inaccessible to such imaging. If one had a gene that selectively responds to such pulses, one could stain for such a gene to perform higher-throughput and *in vivo* fixed assays. Because no genes with selective pulse detection have been identified/characterized, we decided to engineer such a gene.

To build such a synthetic gene, we first had to identify networks that could enable robust pulse detection. We decided to focus on the feed-forward loop (FFL) network architecture class. FFLs are a pattern of genes in which an initial input acts an intermediate node and both then go on to act upon the final output node. If both the direct connection and the indirect connection from the stimulus to the output node have the same sign (both activating or both inactivating) the motif is deemed coherent; in the opposite scenario it is considered incoherent[8,9]. In the past, detailed characterization of these network motifs has revealed their wide range of filtering capabilities. For instance, the coherent FFL type 1 has been shown to enable persistence detection – sustained but not pulsatile inputs lead to expression of the final output[10]. Incoherent FFLs have been implicated in enabling fold-change detection, specifically in the case of WNT signaling[11]. Finally, it has been shown that many of the incoherent FFLs can respond to pulses, however their selectivity over sustained on or off signals has yet to be determined[12].

##### 1.2 Computational screen using AND logic

To determine which FFL architecture could implement pulse detection, we decided to first assay all eight of the possible combinations for a FFL with three connections using both AND and OR logic for the final output node. To build these models we started from a simple set of two equations in which the intermediate node (x1) is made in proportion to the amount of input and the output node (x2) is made by integrating the amounts of x1 and input. As an example, below is the derivation for Circuit 2 (**Figure 1B**):

$$\frac{dx_1}{dt} = V_1 \cdot \frac{C_1^{n_1}}{C_1^{n_1}+u^{n_1}} - k_1 \cdot x_1 \qquad (1)$$

$$\frac{dx_2}{dt} = V_2 \cdot \frac{x_1^{n_2}}{x_1^{n_2}+C_2^{n_2}} \cdot \frac{C_3^{n_3}}{u^{n_3}+C_3^{n_3}} - k_2 \cdot x_2 \qquad (2)$$

In Eq.1, *u* represents the signaling input and has values of either 0 uM when the input is off, and a value of 10 uM when the input is on. Species *x1* is produced in a manner proportional to the parameter $V_1$, and in proportion to one Hill function. In this Hill function, *u* acts as an inhibitor and $C_1$ is the activation coefficient. As *u* surpasses $C_1$ the fraction approaches 0 making the production term go to 0. On the other hand, when *u* is less than $C_1$, the Hill function is close to 1, making the production term a positive number. The Hill coefficient $n_1$ govern the steepness of this response. Species $x_1$ degrades at a rate proportional to itself and parameter $k_1$. The structure of Eq. 2 resembles the structure of Eq. 1, except there are now two hill functions integrated in AND logic. The activator in the first Hill function is $x_1$ and the repressor in the second Hill function is *u*.

Next, we nondimensionalize Eq. 1-2 using the identities in Eq. 3-5 to focus in on the characteristic dynamics of the READer system.

$$\widetilde{x_1} = \frac{x_1}{V_1/k_1} \tag{3}$$

$$\widetilde{x_2} = \frac{x_2}{V_2/k_2} \tag{4}$$

$$\tilde{u} = \frac{u}{C_1} \tag{5}$$

We also define the following nondimensional parameters:

$$\alpha = \frac{C_3}{C_1} \tag{6}$$

$$\beta = \frac{C_2}{V_2/k_2} \tag{7}$$

Inserting these nondimensional identities and parameters into Eq. 1-2 produces the equations used for Circuit 2 in the computational screen (**Figure 1D, Figure S3**):

$$\frac{d\widetilde{x_1}}{dt} = k_1\left(\frac{\alpha^{n_1}}{u^{n_1} + \alpha^{n_1}} - \widetilde{x_1}\right) \tag{8}$$

$$\frac{d\widetilde{x_2}}{dt} = k_2\left(\frac{\widetilde{x_1}^{n_2}}{\widetilde{x_1}^{n_2} + \beta^{n_2}} \cdot \frac{\alpha^{n_3}}{\tilde{u}^{n_3} + \alpha^{n_3}} - \widetilde{x_2}\right) \tag{9}$$

From this derivation one can appreciate that a general form for the FFL loops is as follows:

$$\frac{d\widetilde{x_1}}{dt} = k_1\big(f_{+/-}(u) - \widetilde{x_1}\big) \tag{10}$$

$$\frac{d\widetilde{x_2}}{dt} = k_2\big(g_{+/-}(\widetilde{x_1}) \cdot h_{+/-}(u) - \widetilde{x_2}\big) \tag{11}$$

In these equations, the positive (activating) form for the hill functions *f*, *g,* or *h* uses the form:

$$\frac{x^n}{x^n + \beta^n} \tag{12}$$

And the negative (repressive) form of the hill function uses the form:

$$\frac{\beta^n}{x^n + \beta^n} \tag{13}$$

For equations 12-13, in the case of functions *f* and *h* in equations 10-11, $u$ would take the place of $x$ and $\alpha$ would take the place of $\beta$.

In the following table, we delineate all of the equations used in the computational screen for each incoherent feedforward circuit with AND logic:

| | |
|---|---|
| Circuit 1 | $\frac{d\widetilde{x_1}}{dt} = k_1\left(\frac{u^{n_1}}{u^{n_1}+1} - \widetilde{x_1}\right)$ <br> $\frac{d\widetilde{x_2}}{dt} = k_2\left(\frac{\widetilde{x_1}^{n_2}}{\widetilde{x_1}^{n_2}+\beta^{n_2}} \cdot \frac{u^{n_3}}{u^{n_3}+\alpha^{n_3}} - \widetilde{x_2}\right)$ |
| Circuit 2 | $\frac{d\widetilde{x_1}}{dt} = k_1\left(\frac{\alpha^{n_1}}{u^{n_1}+\alpha^{n_1}} - \widetilde{x_1}\right)$ <br> $\frac{d\widetilde{x_2}}{dt} = k_2\left(\frac{\widetilde{x_1}^{n_2}}{\widetilde{x_1}^{n_2}+\beta^{n_2}} \cdot \frac{\alpha^{n_3}}{u^{n_3}+\alpha^{n_3}} - \widetilde{x_2}\right)$ |
| Circuit 3 | $\frac{d\widetilde{x_1}}{dt} = k_1\left(\frac{u^{n_1}}{u^{n_1}+1} - \widetilde{x_1}\right)$ <br> $\frac{d\widetilde{x_2}}{dt} = k_2\left(\frac{\beta^{n_2}}{\widetilde{x_1}^{n_2}+\beta^{n_2}} \cdot \frac{\alpha^{n_3}}{u^{n_3}+\alpha^{n_3}} - \widetilde{x_2}\right)$ |
| Circuit 4 | $\frac{d\widetilde{x_1}}{dt} = k_1\left(\frac{\alpha^{n_1}}{u^{n_1}+\alpha^{n_1}} - \widetilde{x_1}\right)$ <br> $\frac{d\widetilde{x_2}}{dt} = k_2\left(\frac{\widetilde{x_1}^{n_2}}{\widetilde{x_1}^{n_2}+\beta^{n_2}} \cdot \frac{\alpha^{n_3}}{u^{n_3}+\alpha^{n_3}} - \widetilde{x_2}\right)$ |
| Circuit 5 | $\frac{d\widetilde{x_1}}{dt} = k_1\left(\frac{u^{n_1}}{u^{n_1}+1} - \widetilde{x_1}\right)$ <br> $\frac{d\widetilde{x_2}}{dt} = k_2\left(\frac{\widetilde{x_1}^{n_2}}{\widetilde{x_1}^{n_2}+\beta^{n_2}} \cdot \frac{\alpha^{n_3}}{u^{n_3}+\alpha^{n_3}} - \widetilde{x_2}\right)$ |

| | |
|---|---|
| Circuit 6 | $$\frac{d\widetilde{x_1}}{dt} = k_1\left(\frac{\alpha^{n_1}}{u^{n_1} + \alpha^{n_1}} - \widetilde{x_1}\right)$$ $$\frac{d\widetilde{x_2}}{dt} = k_2\left(\frac{\beta^{n_2}}{\widetilde{x_1}^{n_2} + \beta^{n_2}} \cdot \frac{\alpha^{n_3}}{u^{n_3} + \alpha^{n_3}} - \widetilde{x_2}\right)$$ |
| Circuit 7 | $$\frac{d\widetilde{x_1}}{dt} = k_1\left(\frac{u^{n_1}}{u^{n_1} + 1} - \widetilde{x_1}\right)$$ $$\frac{d\widetilde{x_2}}{dt} = k_2\left(\frac{\beta^{n_2}}{\widetilde{x_1}^{n_2} + \beta^{n_2}} \cdot \frac{u^{n_3}}{u^{n_3} + \alpha^{n_3}} - \widetilde{x_2}\right)$$ |
| Circuit 8 | $$\frac{d\widetilde{x_1}}{dt} = k_1\left(\frac{\alpha^{n_1}}{u^{n_1} + \alpha^{n_1}} - \widetilde{x_1}\right)$$ $$\frac{d\widetilde{x_2}}{dt} = k_2\left(\frac{\widetilde{x_1}^{n_2}}{\widetilde{x_1}^{n_2} + \beta^{n_2}} \cdot \frac{u^{n_3}}{u^{n_3} + \alpha^{n_3}} - \widetilde{x_2}\right)$$ |

To perform the computational screen, we applied three separate inputs: 0.00001 (constant off), 4 (pulsed), and 40 simulation time units (constant on) to a randomly chosen set of parameters for each circuit for a total simulation time of 40 time unites. The area under the curve (AUC) of the $x_2$ curve for each simulation was used to calculate the output under each input condition. By computing the ratio of the AUC of pulsed to those of the other constant inputs, we could identify simulations that enabled selective pulse detection. This was done 10,000 times for each circuit (**Figure 1C-D**). From this analysis, only Circuit 7 showed any robust level of pulse detection, with >90% of simulations resulting in greater output in the pulsed condition compared to the constant on and off inputs.

To ensure that our screen recapitulated known features of FFL, we looked at each individual circuit under the same parameter values. From this analysis, it was clear that Circuit 7 did provide pulse detection while the other circuits did not (**Figure 1E, Figure S3**). We also saw that

Circuit 1, also known as Coherent FFL type 1, responded most in the sustained constant on input case (**Figure S3A**). This is in line with previous literature that suggests that this network motif enables persistence detection[10].

**1.3 Computational screen using OR logic**

In the previous computational screen, we computed the final output node (x2) using AND logic. By this we mean that both input and x1 need to be positively acting on x2 at the same time to produce x2; if either input or x1 is a repressor, then their absence is require for the production of x2. We next wanted to perform the same screen except using OR logic for x2 when it integrates the levels of both x1 and input. To do this, we changed the form of equation 11 to the following:

$$\frac{d\widetilde{x_2}}{dt} = k_2\big(1 - (1 - g_{+/-}(\widetilde{x_1})) \cdot (1 - h_{+/-}(u)) - \widetilde{x_2}\big) \tag{14}$$

In this setup, let us say that both *g* and *h* are activating functions (eq. 12). In the case when either $x_1$ or $u$ is greater than their respective threshold value, then, the inner (1- $g_{+/-}(\widetilde{x_1})$) or $(1 - h_{+/-}(u))$ go to 0 making the production term 1. Only in the case that both g($x_1$) and h($u$) are approaching 0 does the production term go to 0. Using this logic, we arrive at the following 8 sets of equations for incoherent feedforward networks using OR logic:

| | |
|---|---|
| Circuit 1 | $\frac{d\widetilde{x_1}}{dt} = k_1\left(\frac{u^{n_1}}{u^{n_1}+1} - \widetilde{x_1}\right)$ <br> $\frac{d\widetilde{x_2}}{dt} = k_2\left(1 - (1 - \frac{\widetilde{x_1}^{n_2}}{\widetilde{x_1}^{n_2}+\beta^{n_2}}) \cdot (1 - \frac{u^{n_3}}{u^{n_3}+\alpha^{n_3}}) - \widetilde{x_2}\right)$ |

| | |
|---|---|
| Circuit 2 | $\frac{d\widetilde{x_1}}{dt} = k_1\left(\frac{\alpha^{n_1}}{u^{n_1} + \alpha^{n_1}} - \widetilde{x_1}\right)$ <br> $\frac{d\widetilde{x_2}}{dt} = k_2\left(1 - (1 - \frac{\widetilde{x_1}^{n_2}}{\widetilde{x_1}^{n_2} + \beta^{n_2}}) \cdot (1 - \frac{\alpha^{n_3}}{u^{n_3} + \alpha^{n_3}}) - \widetilde{x_2}\right)$ |
| Circuit 3 | $\frac{d\widetilde{x_1}}{dt} = k_1\left(\frac{u^{n_1}}{u^{n_1} + 1} - \widetilde{x_1}\right)$ <br> $\frac{d\widetilde{x_2}}{dt} = k_2\left(1 - (1 - \frac{\beta^{n_2}}{\widetilde{x_1}^{n_2} + \beta^{n_2}}) \cdot (1 - \frac{\alpha^{n_3}}{u^{n_3} + \alpha^{n_3}}) - \widetilde{x_2}\right)$ |
| Circuit 4 | $\frac{d\widetilde{x_1}}{dt} = k_1\left(\frac{\alpha^{n_1}}{u^{n_1} + \alpha^{n_1}} - \widetilde{x_1}\right)$ <br> $\frac{d\widetilde{x_2}}{dt} = k_2\left(1 - (1 - \frac{\widetilde{x_1}^{n_2}}{\widetilde{x_1}^{n_2} + \beta^{n_2}}) \cdot (1 - \frac{\alpha^{n_3}}{u^{n_3} + \alpha^{n_3}}) - \widetilde{x_2}\right)$ |
| Circuit 5 | $\frac{d\widetilde{x_1}}{dt} = k_1\left(\frac{u^{n_1}}{u^{n_1} + 1} - \widetilde{x_1}\right)$ <br> $\frac{d\widetilde{x_2}}{dt} = k_2\left(1 - (1 - \frac{\widetilde{x_1}^{n_2}}{\widetilde{x_1}^{n_2} + \beta^{n_2}}) \cdot (1 - \frac{\alpha^{n_3}}{u^{n_3} + \alpha^{n_3}}) - \widetilde{x_2}\right)$ |
| Circuit 6 | $\frac{d\widetilde{x_1}}{dt} = k_1\left(\frac{\alpha^{n_1}}{u^{n_1} + \alpha^{n_1}} - \widetilde{x_1}\right)$ <br> $\frac{d\widetilde{x_2}}{dt} = k_2\left(1 - (1 - \frac{\beta^{n_2}}{\widetilde{x_1}^{n_2} + \beta^{n_2}}) \cdot (1 - \frac{\alpha^{n_3}}{u^{n_3} + \alpha^{n_3}}) - \widetilde{x_2}\right)$ |
| Circuit 7 | $\frac{d\widetilde{x_1}}{dt} = k_1\left(\frac{u^{n_1}}{u^{n_1} + 1} - \widetilde{x_1}\right)$ <br> $\frac{d\widetilde{x_2}}{dt} = k_2\left(1 - (1 - \frac{\beta^{n_2}}{\widetilde{x_1}^{n_2} + \beta^{n_2}}) \cdot (1 - \frac{u^{n_3}}{u^{n_3} + \alpha^{n_3}}) - \widetilde{x_2}\right)$ |
| Circuit 8 | $\frac{d\widetilde{x_1}}{dt} = k_1\left(\frac{\alpha^{n_1}}{u^{n_1} + \alpha^{n_1}} - \widetilde{x_1}\right)$ <br> $\frac{d\widetilde{x_2}}{dt} = k_2\left(1 - (1 - \frac{\widetilde{x_1}^{n_2}}{\widetilde{x_1}^{n_2} + \beta^{n_2}}) \cdot (1 - \frac{u^{n_3}}{u^{n_3} + \alpha^{n_3}}) - \widetilde{x_2}\right)$ |

We then perform the same computational screen that we performed for the AND gate logic. From this analysis, it was clear that none of the OR gate logics performed pulse detection as few simulations ever landed in the upper right hand quadrant (**Figure S2A**). However, one circuit seemed to display pulse specific repression – Circuit 5 (**Figure S2B**). Interestingly, this circuit is the logical inverse of Circuit 7 with AND logic. Circuit 7 with AND logic can be represented as:

$$x_2 = NOT\ u\ AND\ x_1 \qquad (15)$$

If $x_2$ is pulse detection and we are looking for pulse specific repression, we are essentially searching for (NOT $x_2$). When we perform this NOT operation over eq. 15, we get:

$$\widetilde{x_2} = NOT\ x_2 = u\ OR\ NOT\ x_1 \qquad (16)$$

The logical operation represented by eq. 16 is the one performed in Circuit 5 with OR logic.

**1.4 Conclusion**

Overall, we have successfully performed a computational screen over all FFL networks using both AND and OR logic for the final output node to search for pulse detecting circuits. Validating our approach, Circuit 1 (which has previously been identified as a persistence detection circuit) robustly responded to sustained signals more strongly than pulsed and off inputs. Using this screen, we identified one circuit topology – Circuit 7 – which provides pulse detection. Interestingly, our OR gate screen revealed that the logical inverse of Circuit 7 with AND gate logic, Circuit 5, allowed for pulse-specific repression, further validating the approach. This work, in conjunction with studies[10,13–15], demonstrates the utility of computational screens for identifying biological networks that enable specific filtering capabilities.

**2. Characterization of 3-node network that enables band-pass filtering**

#### 2.1. Goal: Understand and experimentally tune parameters controlling band-pass filtering

To our knowledge, this is the first network motif that has been described to possess such selective dynamic band-pass filtering capabilities at the protein level, providing the first insight into how cells may interpret different frequencies. However, how can different genes respond to different frequencies? What parameters do cells alter to change the selectivity and placement of the optimal response pulse? To address these questions, we used both mathematical modeling and experiments to understand what affects properties of the band-pass such as peak position, peak height and band-pass width (selectivity). To computationally assay the system, we took our model (**Figure 2H**) that has three equations for the three species (transcription of KTR-Gal4, GFP and negative regulator) with its 11 parameters and varied each parameter independently 100-fold down and up from the values previous used that seemed to recapitulate experimental data (**Figure 2I**). For each parameter set, we simulated a band-pass experiment in which we initiate pulses of different lengths and then wait a fixed amount of simulated time. Each resulting band-pass was then analyzed for the peak position, peak height, and width (selectivity). From this analysis, it was clear that there were 3 key parameters that affected these features: the timescale on which the KTR-Gal4 is made, the timescale that the negative regulator is made, and the affinity that Gal4 protein has for the GFP promoter (**Figure S12**). By simply altering these three parameters, we could define band-pass curves of similar selectivity and peak height, for different peak positions (**Figure S13**).

#### 2.2: Mathematical model for parameter scan

Equations 17-19 describe the dynamics of the READer system. For simplicity, we do not represent every component of the READer system with an equation. Instead, we choose the minimal number of equations necessary to reproduce the most salient features of the READer system dynamics: pulse detection and band-pass filtering. Species $x_1$ can be thought of as the transcription of KTR-Gal4, both in the active and inactive form. Species $x_2$ can be thought of as the amount of nuclear KTR-Gal4 protein. Species $x_3$ can be thought of as the amount of ERK negative regulator in cells. At t = 0 min, $x_1 = x_2 = x_3 = 0$ uM.

$$\frac{dx_1}{dt} = V_1 \cdot \frac{u^{n_1}}{C_1^{n_1} + u^{n_1}} \cdot \frac{C_2^{n_2}}{x_3^{n_2} + C_2^{n_2}} - k_1 \cdot x_1 \quad (17)$$

$$\frac{dx_2}{dt} = V_2 \cdot \frac{x_1^{n_3}}{x_1^{n_3} + C_3^{n_3}} \cdot \frac{C_4^{n_4}}{u^{n_4} + C_4^{n_4}} - k_2 \cdot x_2 \quad (18)$$

$$\frac{dx_3}{dt} = V_3 \cdot \frac{x_1^{n_5}}{x_1^{n_5} + C_5^{n_5}} - k_3 \cdot x_3 \quad (19)$$

In Eq. 17, $u$ represents the ERK signaling input, which is on in the presence of growth factors or optogenetic stimulation and off when growth factors and optogenetic stimulation are absent. When off, $u$ has a value of 0 μM, and when on $u$ has a value of 10 μM. Species $x_1$ is produced in a manner proportional to the parameter $V_1$, and in proportion to two Hill functions. In the first Hill function, $u$ acts as an activator and $C_1$ is the activation coefficient. In the second Hill function, $x_3$ acts as a repressor and $C_2$ is the repression coefficient. The Hill coefficients $n_1$ and $n_2$ govern the steepness of the first and second Hill functions, respectively. Species $x_1$ degrades at a rate proportional to itself and parameter $k_1$. The structure of Eq. 18 resembles the structure of Eq. 17, except the activator in the first Hill function is $x_1$ and the repressor in the second Hill function is $u$. Eq. 19 only contains a single Hill function with activator $x_1$.

We nondimensionalize Eq. 17-19 using the identities in Eq. 20-23 to focus in on the characteristic dynamics of the READer system.

$$\widetilde{x_1} = \frac{x_1}{V_1/_{k_1}} \tag{20}$$

$$\widetilde{x_2} = \frac{x_2}{V_2/_{k_2}} \tag{21}$$

$$\widetilde{x_3} = \frac{x_3}{V_3/_{k_3}} \tag{22}$$

$$\tilde{u} = \frac{u}{C_1} \tag{23}$$

We also define the following nondimensional parameters:

$$\alpha = \frac{C_4}{C_1} \tag{24}$$

$$\beta_1 = \frac{C_2}{V_3/_{k_3}} \tag{25}$$

$$\beta_2 = \frac{C_3}{V_1/_{k_1}} \tag{26}$$

$$\beta_3 = \frac{C_5}{V_1/_{k_1}} \tag{27}$$

Inserting Eq.20-27 nondimensional identities and parameters into Eq. 17-19 produces the nondimensionalized equations used in the main text (**Figure 2G**):

$$\frac{d\widetilde{x_1}}{dt} = k_1\left(\frac{\tilde{u}^{n_1}}{1+\tilde{u}^{n_1}} \cdot \frac{\beta_1^{n_2}}{\widetilde{x_3}^{n_2}+\beta_1^{n_2}} - \widetilde{x_1}\right) \tag{28}$$

$$\frac{d\widetilde{x_2}}{dt} = k_2 \left( \frac{\widetilde{x_1}^{n_3}}{\widetilde{x_1}^{n_3} + \beta_2^{n_3}} \cdot \frac{\alpha^{n_4}}{\tilde{u}^{n_4} + \alpha^{n_4}} - \widetilde{x_2} \right) \tag{29}$$

$$\frac{d\widetilde{x_3}}{dt} = k_3 \left( \frac{\widetilde{x_1}^{n_5}}{\widetilde{x_1}^{n_5} + \beta_3^{n_5}} - \widetilde{x_3} \right) \tag{30}$$

As discussed in the main text, the READer system serves as a bandpass filter that discriminates among different durations of ERK signaling. Using the set of equations from 28-30, we can reproduce all of the important dynamic filtering capabilities: (1) detection of a single short pulse of input (**Figure 2D-G**), (2) ignoring of long pulses (**Figure 2G**), and (3) band-pass filtering depending on Eq. 30 (negative feedback), meaning that both very short pulses and very long pulses do not result in maximum output and that an intermediate pulse results in the optimal output of the system (**Figure 2I**).

**Table 1. Initial parameters for dynamic filtering computational screen.**

| Parameter | Value [units] |
|---|---|
| $n_1$ | 1 |
| $n_2$ | 5 |
| $n_3$ | 5 |
| $n_4$ | 5 |
| $n_5$ | 5 |
| $k_1$ | 0.1 [1/min] |
| $k_2$ | 0.02 [1/min] |
| $k_3$ | 0.025 [1/min] |

| | |
|---|---|
| $\beta_1$ | 0.2 |
| $\beta_2$ | 0.1 |
| $\beta_3$ | 0.01 |
| [illegible] | 0.5 |

For the model, the values above were picked by hand to reflect the time scales of KTR and GFP production and degradation observed in the time course data (**Figure S5**). The variable $\widetilde{x_2}$ corresponds to the amount GFP transcription induced by nuclear KTR-Gal4. We can generate a computational bandpass curve from Equations 28-30 by plotting the area under the curve of the $\widetilde{x_2}$ trace for different values of pulse length $\tau$ where:

$$\tilde{u} = 10\ \mu M, (0 \leq t \leq \tau) \quad (31)$$
$$\tilde{u} = 0\ \mu M, (t > \tau)$$

The total simulation time for each $\tau$ is $\tau$ +100 time units to allow for GFP to be made. The bandpass curve for the parameter values listed in Table 1 is centered around (i.e. $\widetilde{x_2}$ has the highest value for) an ERK signaling duration of t = 41.8 min.

**2.3: Parameter scan for variables that control bandpass filtering**

To begin to understand what controls bandpass filtering we decide to perform a parameter scan for each parameter in eq. 28- 30. We sought to better under how changing parameter values changes the bandpass curve's peak location, peak amplitude, and selectivity. A higher width corresponds to a lower selectivity. Peak location is the pulse length $\tau$ that allows for the highest

GFP and peak amplitude is the resulting GFP from this optimal pulse length. We define the selectivity of the bandpass as the width of the curve at half the maximum $\widetilde{x_2}$ amplitude for each bandpass curve. The results of this analysis would shed light on how the physical properties of the molecules that compose READer and the native ERK-dependent processes it interacts with relate to the dynamic filtering features of READer.

The parameters in **Table 1** relate to physical properties of the molecules that READer interacts with and is composed of. Parameters $k_1$, $k_2$, and $k_3$ dictate the time scale of KTR-Gal4, GFP, and ERK negative regulator production, respectively, with low k values resulting in slow production and high k values resulting in fast production. In Equations 17-19, $k_1$, $k_2$, and $k_3$ appear as the parameters governing the degradation of $\widetilde{x_1}$, $\widetilde{x_2}$, and $\widetilde{x_3}$, respectively. The value of $k_1$ can be altered through manipulation of the KTR-Gal4 degradation tag, and the value of $k_2$ can be altered through manipulation of the GFP degradation tag. The value of $k_3$ would be difficult to alter, as it corresponds to the degradation rate of the unknown ERK negative regulator native to the cells into which READer has been introduced. In all cases, a high k value would correspond to a protein with fast degradation.

$\beta_1$, $\beta_2$, and $\beta_3$ are the threshold values that help determine whether $\widetilde{x_1}$, $\widetilde{x_2}$, and $\widetilde{x_3}$ are produced, respectively. Significant amounts of KTR-Gal4 can only be produced when the amount of ERK negative regulator is less than $\beta_1$, GFP can only be produced in significant amounts when the amount of KTR-Gal4 exceeds $\beta_2$, and negative regulator can only be produced in significant amounts when the amount of active ERK exceeds $\beta_3$. While $\beta_1$ and $\beta_3$ are properties native to the cells into which READer has been introduced, $\beta_2$ can be altered by strengthening or weakening the UAS sequence that governs GFP production. A stronger UAS would lower the value of $\beta_2$ while a weaker UAS would raise the $\beta_2$ value.

Finally, $\alpha$ is the ratio of the u threshold $C_4$ to the u threshold $C_1$. When $u > C_1$, significant amounts of KTR-Gal4 can be produced. When $u < C_4$, significant amounts of GFP can be produced. Because u can only take on two values, 0 for $u_{off}$ or 10 mM for $u_{on}$, the ratio $\alpha = C_4/C_1$ dictates whether or not a high level of GFP production is possible. If $u_{on} > C_1$ and $C_1 > C_4$, i.e. a $< 1$, KTR-Gal4 can be produced, but it is also excluded from the nucleus and thus cannot induce GFP production while ERK signaling is still present. If $u_{on} > C_1$ and $C_1 < C_4$, i.e. $\alpha > 1$, then GFP can be produced while the ERK signal remains on. For READer to function well as a pulse detector, $\alpha$ must be less than 10. The value of $\alpha$ can be altered by changing the nuclear export sequence (NES) on the KTR-Gal4. A stronger NES would lower the value of $\alpha$ while a weaker NES would raise it.

To better understand how to achieve high-selectivity, high-amplitude bandpass filtering at various ERK signaling durations, we varied each parameter's value from one hundredth its Table 1 value to one hundred times the **Table 1** value. The radar plots show how peak position, peak selectivity, and peak amplitude vary in response to variations in $k_1$, $k_2$, $k_3$, $\beta_1$, $\beta_2$, $\beta_3$, and $\alpha$ (**Figure S12**). The parameters that show the greatest variation in one or more of the bandpass filtering metrics, as shown by the greatest difference along one of the spider plot axes between the yellow (low parameter value) and blue (high parameter value) marker, are $k_1$, $k_3$, and $\beta_2$. Although varying each parameter changes the value of multiple metrics, each parameter controls each metric to a different extent. $k_3$ primarily changes the peak position: increasing $k_3$ moves peak position to a higher ERK signaling duration. Moving the peak position to a higher ERK signaling duration by varying $k_3$ also mildly increases the amplitude and selectivity. To center the bandpass peak around a shorter ERK signaling duration, $k_1$ and $\beta_2$ must be varied to maintain high amplitude and selectivity. Increasing $\beta_2$ does nothing to the peak position, but it increases amplitude and decreases selectivity. Increasing $k_1$ changes all three metrics: the peak position moves to a higher signaling duration, selectivity decreases, and amplitude increases. Notably, only two of the important parameters we identified have values that can be engineered in one direction or another. We predict $k_1$ can be altered by changing the degron tag attached to the KTR-Gal4, while we predict $\beta_2$ can be altered by changing the strength of the UAS sequence to which the KTR-Gal4 binds to produce GFP. $k_3$, however, the parameter that most directly corresponds to the peak position, is dictated by the degradation rate of the unknown ERK negative regulator native to the system.

Because each of these three parameters control the three bandpass metrics to different extents, the values of the parameters can be adjusted in concert to achieve high-selectivity, high-amplitude bandpass filters for any ERK signaling duration. Plotting the parameters values that allowed us to achieve high-selectivity, high-amplitude bandpass filtering for ERK signaling durations of 40, 60, 80, 100, and 120 min, it becomes apparent that a curve can be drawn through 3D parameter space to select for bandpass filters of any duration (**Figure S13**).

**2.4: Experimental tuning of band-pass**

To experimentally tune the READer system, we decided to try and change the degradation of the KTR-Gal4 protein and thus the timescale of the KTR-Gal4 ($k_1$ in our model). Intuitively, if the KTR-Gal4 is degraded faster then fewer of the longer pulse durations will make it to GFP output. To test this, we needed a way to quickly turn on and off signaling on demand in clones that express the reporter GFP gene at equivalent levels. For the first, we used an optogenetic Ras/Erk activator previously created in our lab in which the catalytic domain of SOS is recruited to the membrane *via* the blue-light dimerization system, iLID and SSPB. We took cells expressing this system and infected them with the reporter GFP construct and sorted clones. To ensure that a clone had functional optogenetic Erk activation, we infected clones with ErkKTR-iRFP, placed them on the scope and subjected the cells to cycles of blue light and dark (**Figure S10).** From this analysis we chose one clone that had a functional optogenetic actuator and the reporter.

With this clone in hand we could now test variants of the READer circuit. To increase the degradation of the circuit at the protein level, we either added a modified PEST sequence that decreases the half-life even further than the original PEST sequence. At the transcriptional level we used either the original *TUBULIN* 3' UTR or the one from *FOS*, which has been shown to have a short-lived mRNA. All combinations of these PEST sequences and 3' UTRs were cloned into the READer PiggyBAC construct (**Figure S14A**). These variants were transfected into the test-bed clone and single-cell clones were then derived. When we did a full band-pass experiment with pulse durations ranging from 5 minutes to 12 hours, we found that all 4 READer circuits do implement band-pass filtering (**Figure S14B**). However, when we did a more fine-tuned experiment where we did every pulse duration in increments of 15 minutes from no stimulation to 2 hour pulses, we find that READer constructs harboring the *FOS* 3' UTR have markedly increased selectivity (smaller band-widths) when compared to those that have the longer-lived *TUBULIN* 3' UTR (**Figure S14C-D**).

##### 2.6: Conclusion

Overall, we have successfully developed a minimal mathematical system of equations that recapitulates the bandpass filtering experimentally observed in the READer system. By performing a parameter scan for all of the major components of the system, we find that there are three parameters ($k_1$, $k_3$ and $\beta_2$) that control the selectively, peak position and peak amplitude of the bandpass. Because each of these parameters control more than one of these features, we tune all three simultaneously to derive bandpass curves of specific peak position, height and selectivity. Experimentally we show that by tuning the $k_1$ parameter (degradation of the KTR-Gal4) we are able to change the filtering capabilities of the READer system. This analysis points to the key knobs that nature may alter to specify which pulses may be read out by downstream targets of pulsatile signaling activity. This also provides us a framework with which to develop more synthetic systems to create more tunable and orthogonal dynamic channels.

#### Supplementary Sequences

*fos* 3' UTR

```
gcagtcagagaaggcaaggcagccggcatccagacgtgccactgcccgagctggtgcattacagagagga
gaaacacgtcttccctcgaaggttcccgtcgacctagggaggaccttacctgttcgtgaaacacaccagg
ctgtgggcctcaaggacttgcaagcatccacatctggcctccagtcctcacctcttccagagatgtagca
aaaacaaaacaaaacaaaacaaaaaaccgcatggagtgtgttgttcctagtgacacctgagagctggtag
ttagtagagcatgtgagtcaaggcctggtctgtgtctcttttctctttctccttagttttctcatagcac
taactaatctgttgggttcattattggaattaacctggtgctggattgtatctagtgcagctgattttaa
caatacctactgtgttcctggcaatagcgtgttccaattagaaacgaccaatattaaactaagaaaagat
aggactttattttccagtagatagaaatcaatagctatatccatgtactgtagtccttcagcgtcaatgt
tcattgtcatgttactgatcatgcattgtcgaggtggtctgaatgttctgacattaacagttttccatga
aaacgttttattgtgttttcaatttatttattaagatggattctcagatatttatattttattttatt
tttttctaccctgaggtctttcgacatgtggaaagtgaatttgaatgaaaattttaagcattgtttgct
tattgttccaagacattgtcaataaaagcatttaagttgaa
```

*tubulin* 3' UTR

ttcttaaagcttttactttgagacatcatggaaaacttaagaggtacaacatggagaagacatgatcaca
gaatggaaacagcacagaagcatcagtgcctgcaactaatactggagcagtttgacgacacaggggctc
aaggaatggacttagtactcctctccttcttcctcccctcgccttccacagaatccacaccaacctcct
cataatccttctctagggcagccatgtcctcacgggcctcagagaactcaccctcctccatgccctcacc
cacataccagtgcacaaaggcacgcttggcatacatcagatcaaacttgtgatctaggcgagcccaggcc
tcagcaatggctgtggtgttgctcagcatgcacacagctctctgcaccttggccaggtcaccaccgggta
ccacagtgggaggctggtaattaatgccaaccttgaagccagtggggcaccagtctacaaactggatgct
gcgcttggtcttgatggtggcaatggcagcattgacatctttgggaaccacatcaccacggtatagcagg
cagcaagccatgtatttaccatggcgagggtcacatttcaccatctggttggctggctcaaagcaggcat
tggtgatctctgctacagaaagctgctcatggtaggctttctcagcagagatgacaggggcataagtggc
cagagggaagtggatgcgagggtagggtaccaggttggtctggaattctgtcagatcaacattcagggcc
ccatcaaatctgagggaagcagtgatggaagacacaatctggctaataaggcggttaaggttggtgtagg
ttgggcgctcaatgtcgaggtttctacgacagatgtcatagatggcctcattgtctaccatgaaggcaca
atcagagtgctccagggtggtgtgggtggtgaggatggaattgtagggctcaaccacagcagtggaaacc
tggggggctgggtaaatggagaactccagcttggacttctttccgtaatccacagagagccgctccatca
gcagggaggtgaagccagagccagttcccccaccaaagctgtggaaaaccaagaagccctggagacctgt
gcactggtcagc

KTR-Gal4-VP64 amino acid sequence

MKGRKPRDLELPLSPSLLGGQGPERTPGSGTSSGLQAPGPALSPSKRSGLEDPATPSKKPRTPS
VSSRLERLTLQSSFQFPSTSTRQVEQGRWTVDPVATMKLLSSIEQACDICRLKKLKCSKEKPKC
AKCLKNNWECRYSPKTKRSPLTRAHLTEVESRLERLEQLFLLIFPREDLDMILKMDSLQDIKAL
LTGLFVQDNVNKDAVTDRLASVETDMPLTLRQHRISATSSSEESSNKGQRQLTVSAAAGGSGGS
GGSDALDDFDLDMLGSDALDDFDLDMLGSDALDDFDLDMLGSDALDDFDLDMLGS

#### Supplemental Figures

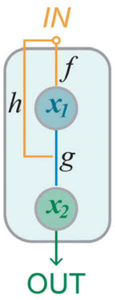


AND gate logic

$$\frac{dx_1}{dt} = k\left(f_{+/-}(u) - x_1\right)$$

$$\frac{dx_2}{dt} = k\left(g_{+/-}(x_1)\,h_{+/-}(u) - x_2\right)$$

positive: $\frac{x^n}{K^n + x^n}$ negative: $\frac{K^n}{K^n + x^n}$

OR gate logic

$$\frac{dx_1}{dt} = k\left(f_{+/-}(u) - x_1\right)$$

$$\frac{dx_2}{dt} = k\left(1 - \left(1 - g_{+/-}(x_1)\right)\left(1 - h_{+/-}(u)\right) - x_2\right)$$

positive: $\frac{x^n}{K^n + x^n}$ negative: $\frac{K^n}{K^n + x^n}$

**Figure S1. Modeling feed-forward loop motifs with AND or OR logic.** Motifs were modeled such that input $u$ acts on the intermediate species $x_1$, which combines with $u$ to regulate the final output node $x_2$. The linkages from one node to the next determine whether $f$, $g$ and $h$ form positive (activating) or negative (repressing) regulatory links. Each activating/repressing connection is based on a Hill function, with $K$ representing the half-maximal concentration of a species $x$ for regulation to occur. The parameter $k$ represents the timescale of changes for each species. The integration of input and $x_1$ is either determined using AND logic or OR logic. With AND logic, $g$ and $h$ are multiplied; thus, output is produced only if the both terms are high. With OR logic, output is produced if either $g$ or $h$ are non-zero through the term 1 – (1-g)(1-h).

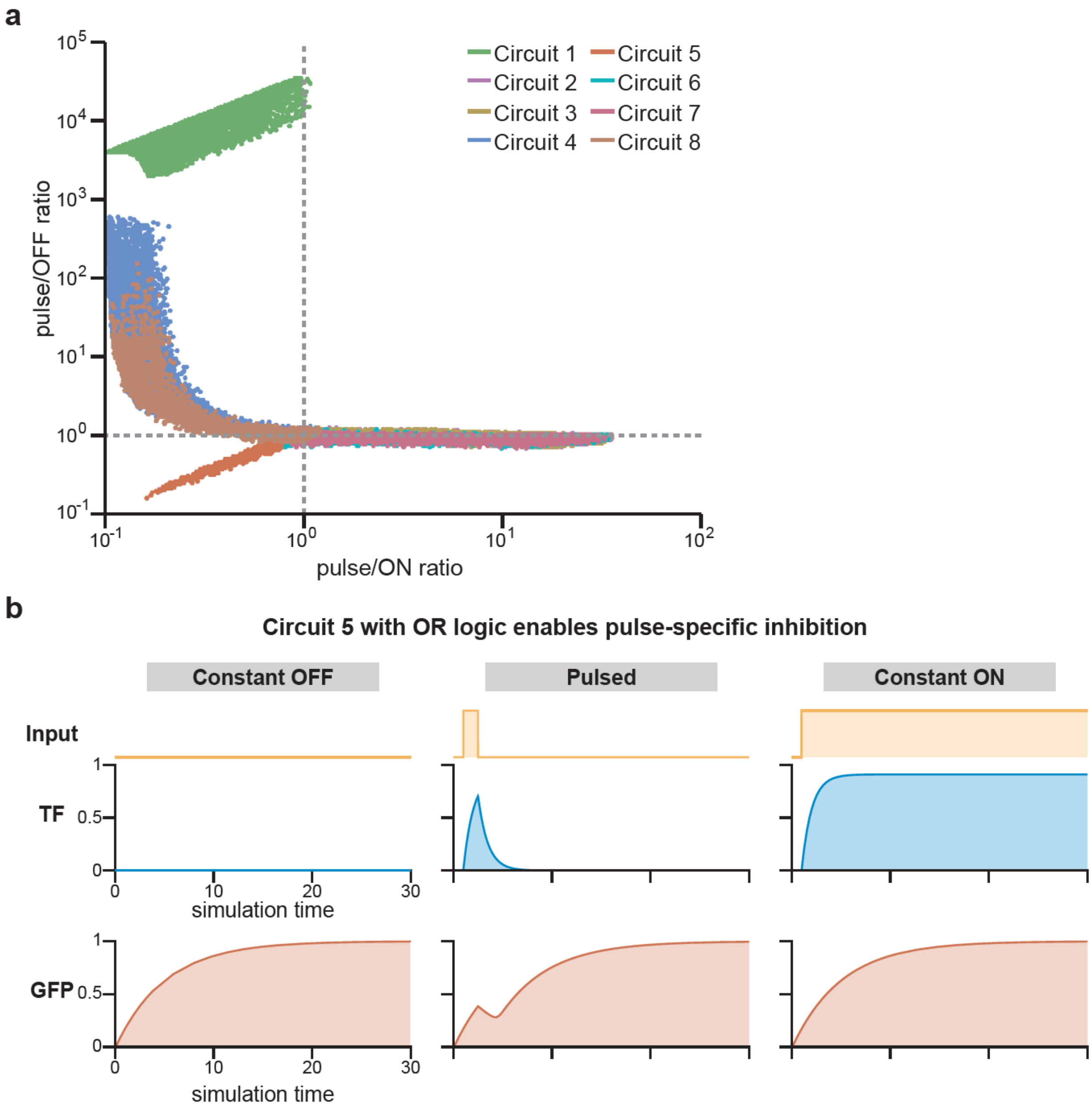


**Figure S2. Computational screen with OR logic reveals pulse-specific inhibition.** (**a**) Plot from computational screen with all 8 circuits (**Figure 1B**) using OR logic. Results of 10,000 simulations per circuit for random choices of parameters. In all cases, the area under the curve of $x_2$ was calculated, and the pulse:OFF and pulse:ON ratios are plotted for each parameter set. (**b**) Representative time course of the pulse-induced repression circuit, Circuit 5, simulated with either constant OFF (left), a pulsed input (middle) or constant ON (right). The pulsed input causes a transient decrease in GFP expression, but no such decrease is observed in constant-off or constant-on cases.

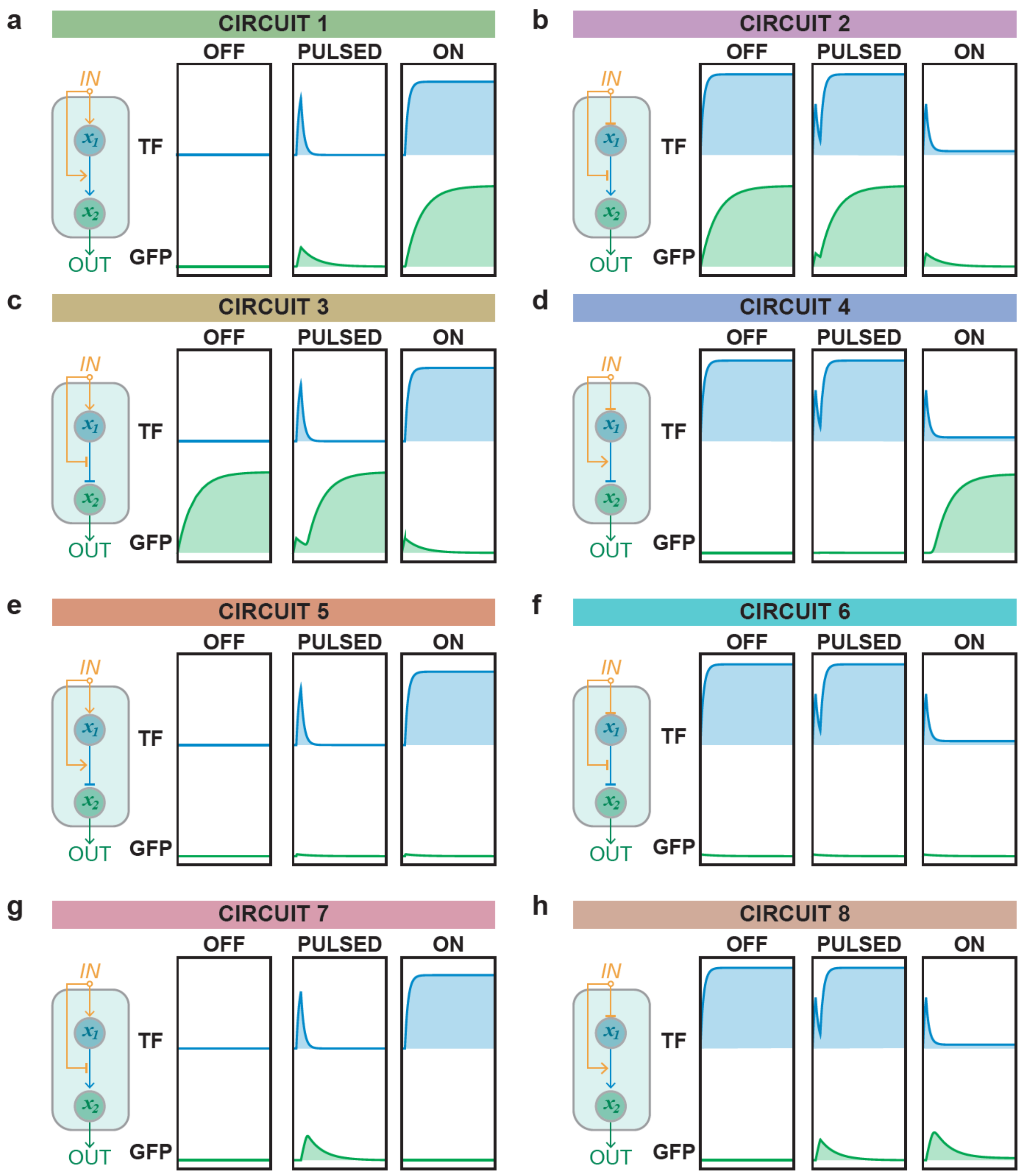


**Figure S3. Representative traces of all 8 circuits using AND logic.** Network topology diagram and representative time course for cases simulated with constant OFF (left), a pulsed input (middle) or constant ON (right) for coherent FFLs [(**a**) circuit 1, (**b**) circuit 2, (**c**) circuit 3, (**d**) circuit 4] and incoherent FFLs [(**e**) circuit 5, (**f**) circuit 6, (**g**) circuit 7, (**h**) circuit 8]. The pulse case for (**g**) circuit 7 are replicated from **Figure 1E.**

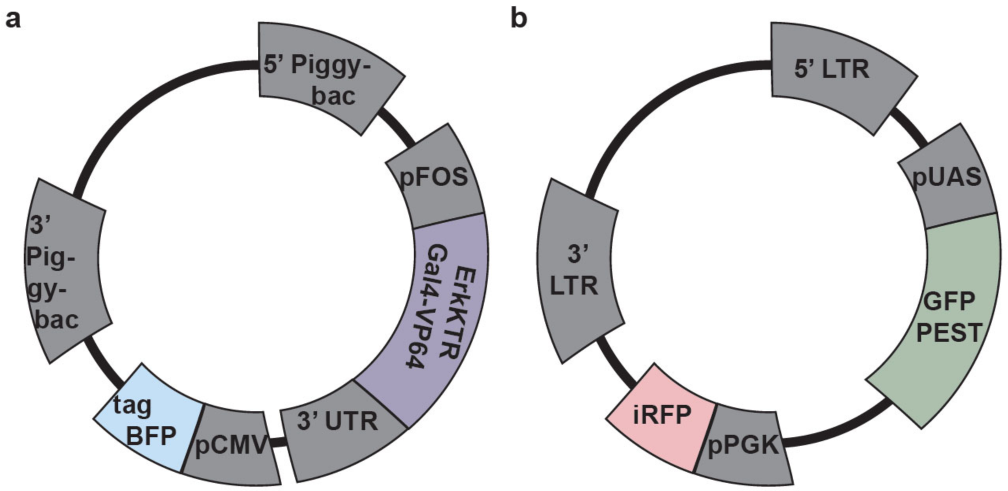


**Figure S4. Maps of key vectors for this study.** (**a**) PiggyBAC vector used to express the dynamics-sensitive transcription factor. This construct is co-transfected along with the PiggyBAC transposase enzyme to enable genomic integration. The CMVp-tagBFP is used as a marker to isolate cells that contain the construct. (**b**) pHR lentivirus vector used to express the UASp-GFP reporter. This construct is co-transfected along with pMD and CMV helper plasmids into Lenti-X 293T cells to generate lentivirus that is then used for genomic integration. The PGKp-iRFP is used to isolate cells that express the construct.

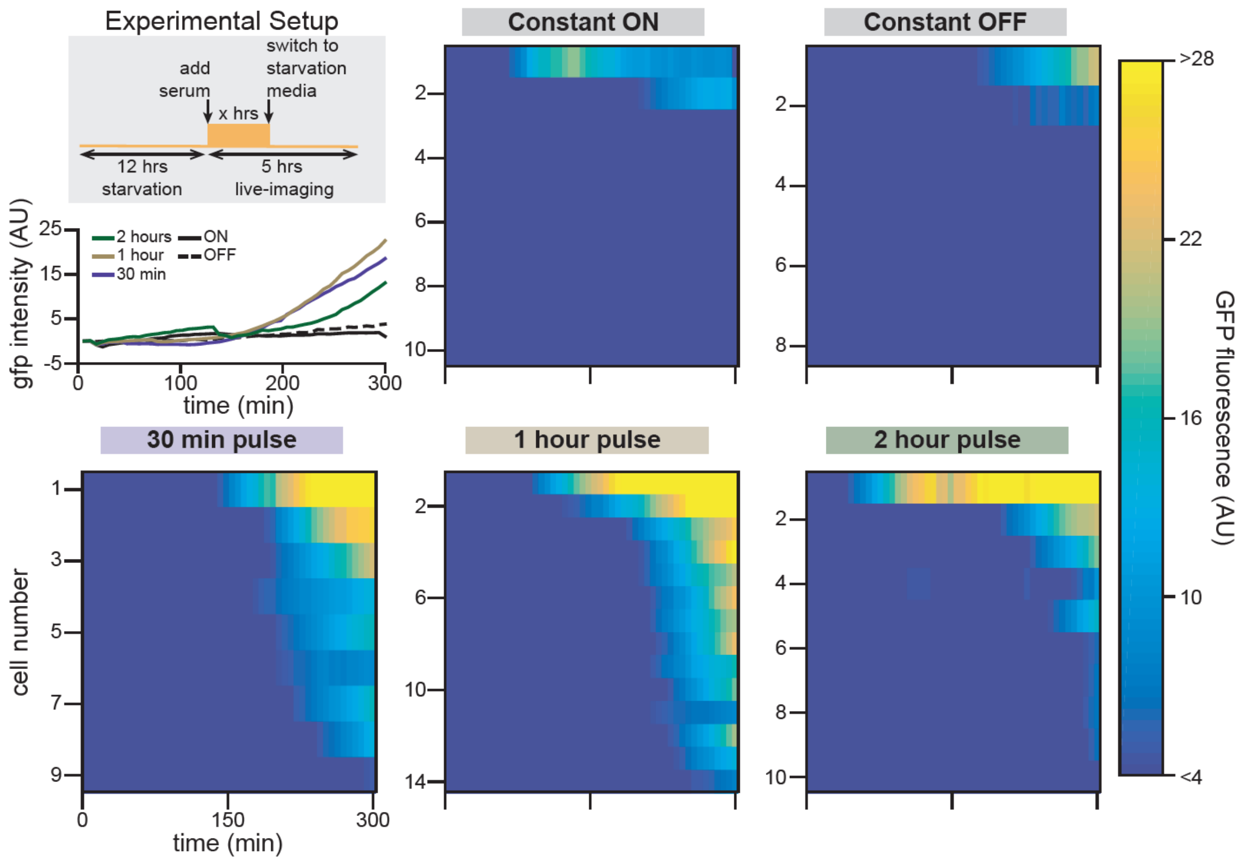


**Figure S5. Confocal imaging reveals that READer acts as a pulse detector.** NIH3T3 READer cells were plated in glass-bottomed 96 plate wells. After 12 hours, cells were switched to growth-factor free media for an additional 12 hours. Curves show mean GFP induction as a function of time for cells in 10% serum (constant ON; n = 10 cells), GF-free media (constant OFF; n=8 cells), a 30-minute serum pulse (n = 9 cells), a 1-hour pulse (n=14 cells) or a 2-hour pulse (n=10 cells). Heatmaps show all single cell traces for the specified condition.

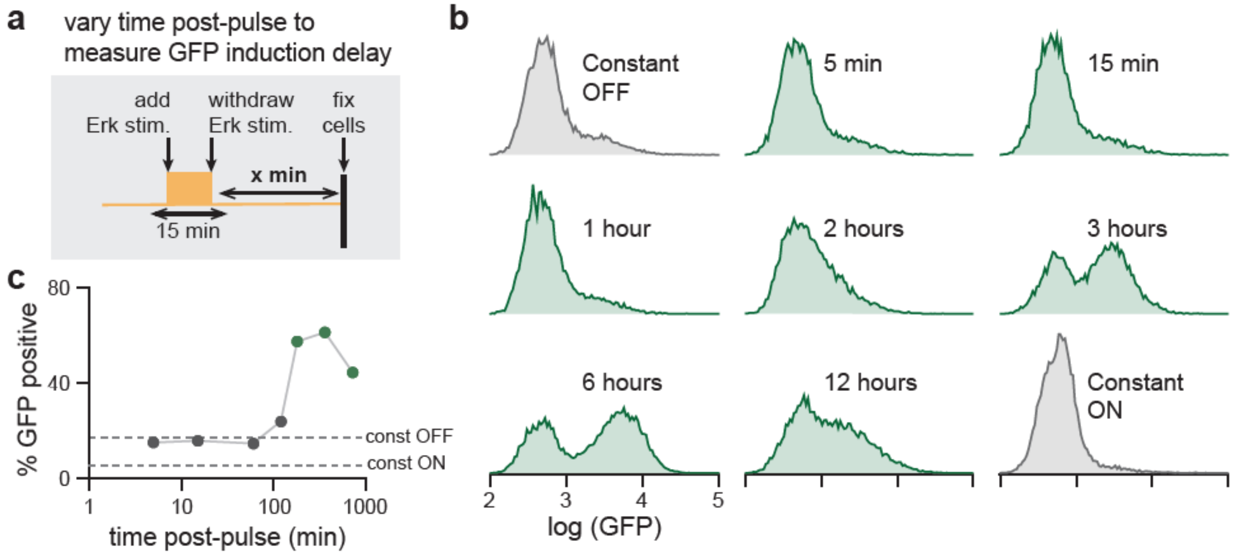


**Figure S6. Quantifying the kinetics of the READer circuit.** (**a-b**) To map how wait-time affects READer circuit output, we subjected cells to a 15 min serum pulse, fixed them at various time-points after the end of the pulse (schematic in **a**) and analyzed by flow cytometry for GFP induction (histograms of GFP expression in **b**). (**c**) Quantification of flow cytometry data in **b** reveals that GFP levels peak ~3 h after the pulse. Points indicate fraction of GFP-positive cells at each time point, with the fraction of GFP-positive cells under constant stimulation conditions shown as dotted lines.

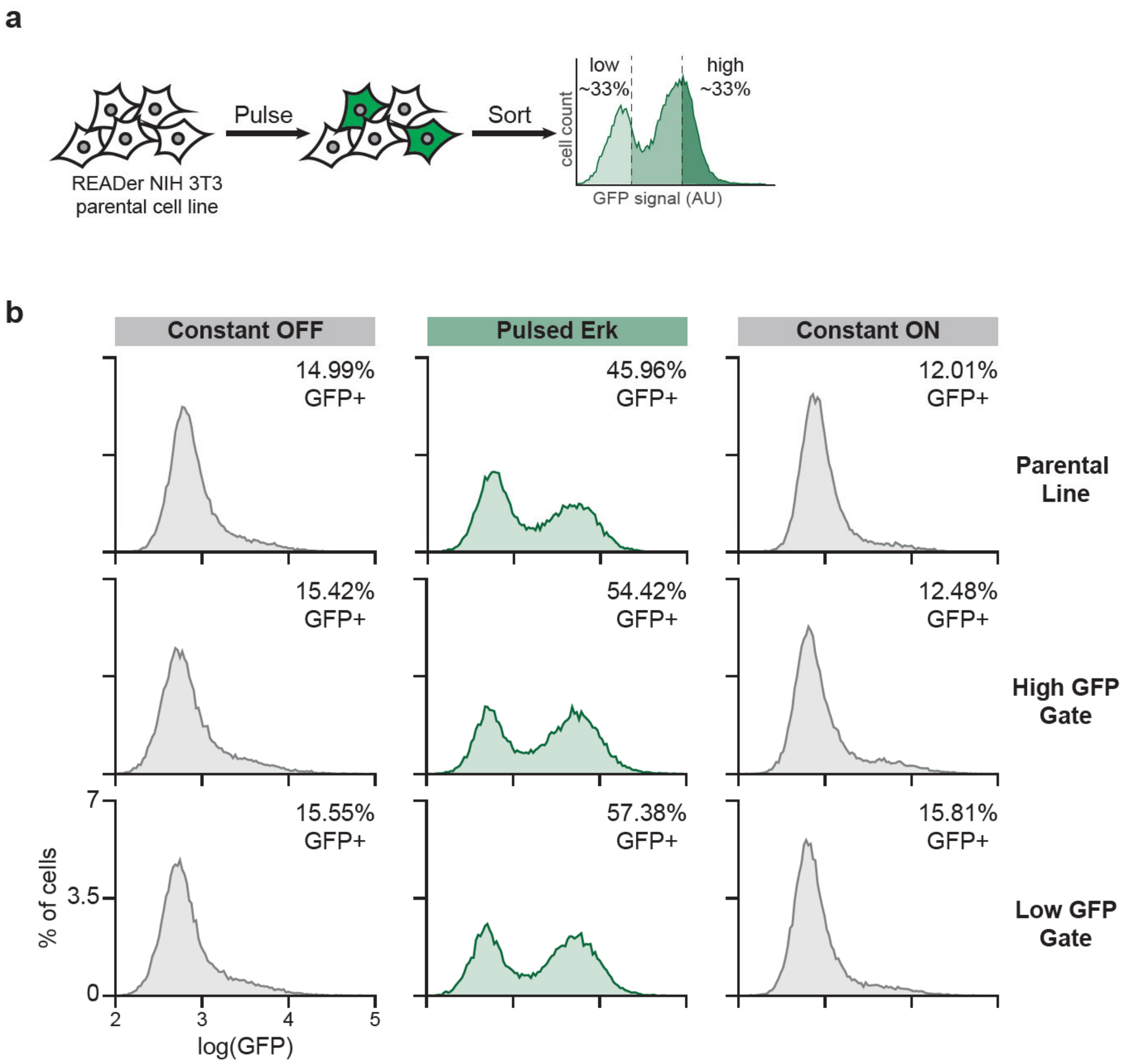


**Figure S7. Non-genetic origin of bimodal READer responses.** (**a**) Schematic of experiment to test whether GFP response is heritable. READer cells were subjected to a 20 min stimulus pulse, incubated for 4 h, and GFP-high and GFP-low subpopulations were sorted. After 1 week, constant and pulsatile stimuli were delivered to parental, GFP-high sorted cells, and GFP-low sorted cells to assess their responses. (**b**) Experimental results. Parental READer cells (top row), GFP-high cells (middle row) and GFP-low cells (bottom row) were analyzed. Cells were starved of growth factors overnight and either received GF-free media (left), a 15 min serum pulse followed by incubation for 6 h (middle) or constant 10% serum (right). Flow cytometry plots along with percent GFP positive cells are shown.

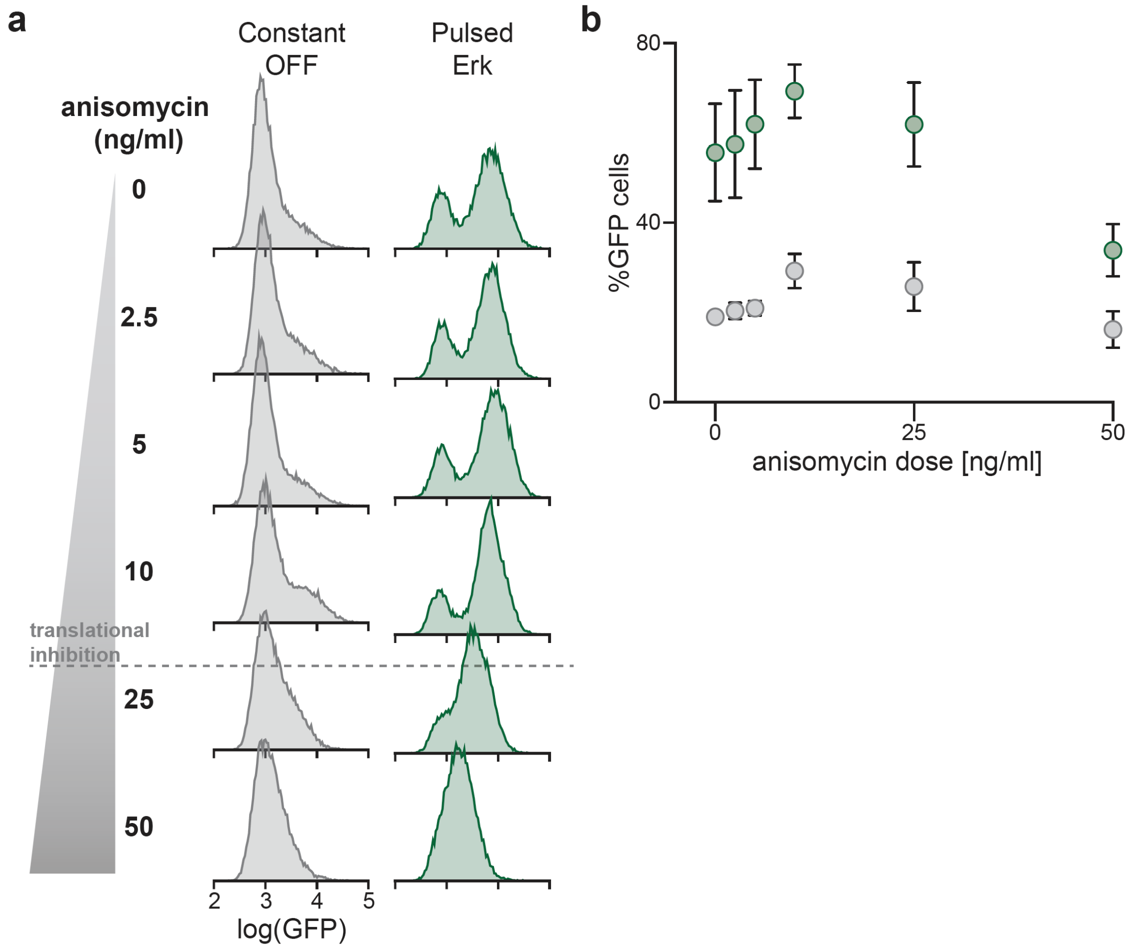


**Figure S8. Anisomycin treatment modulates fraction of responsive READer cells.** (**a**) NIH3T3 READer cells were plated and, after 36 hours, were placed in growth factor free media for 12 hours. Cells then either received GF-free media (Constant OFF) or a pulse of serum (20 minutes on, 4 hours off), each spiked with the indicated concentration of anisomycin before fixation and analysis by flow cytometry. GFP histograms from each condition in one representative experiment are shown. Of note, anisomycin can inhibit translation at concentrations above 10 ng/mL but at lower doses is a potent agonist of p38 and JNK signaling. (**b**) Quantification of GFP positive cells shown in **a** with 2 other replicates for all doses of anisomycin. As expected, high anisomycin doses decrease GFP expression, but intermediate anisomycin doses increase the fraction of GFP+ cells under Erk-pulsing conditions without a corresponding increase in constant-OFF cells.

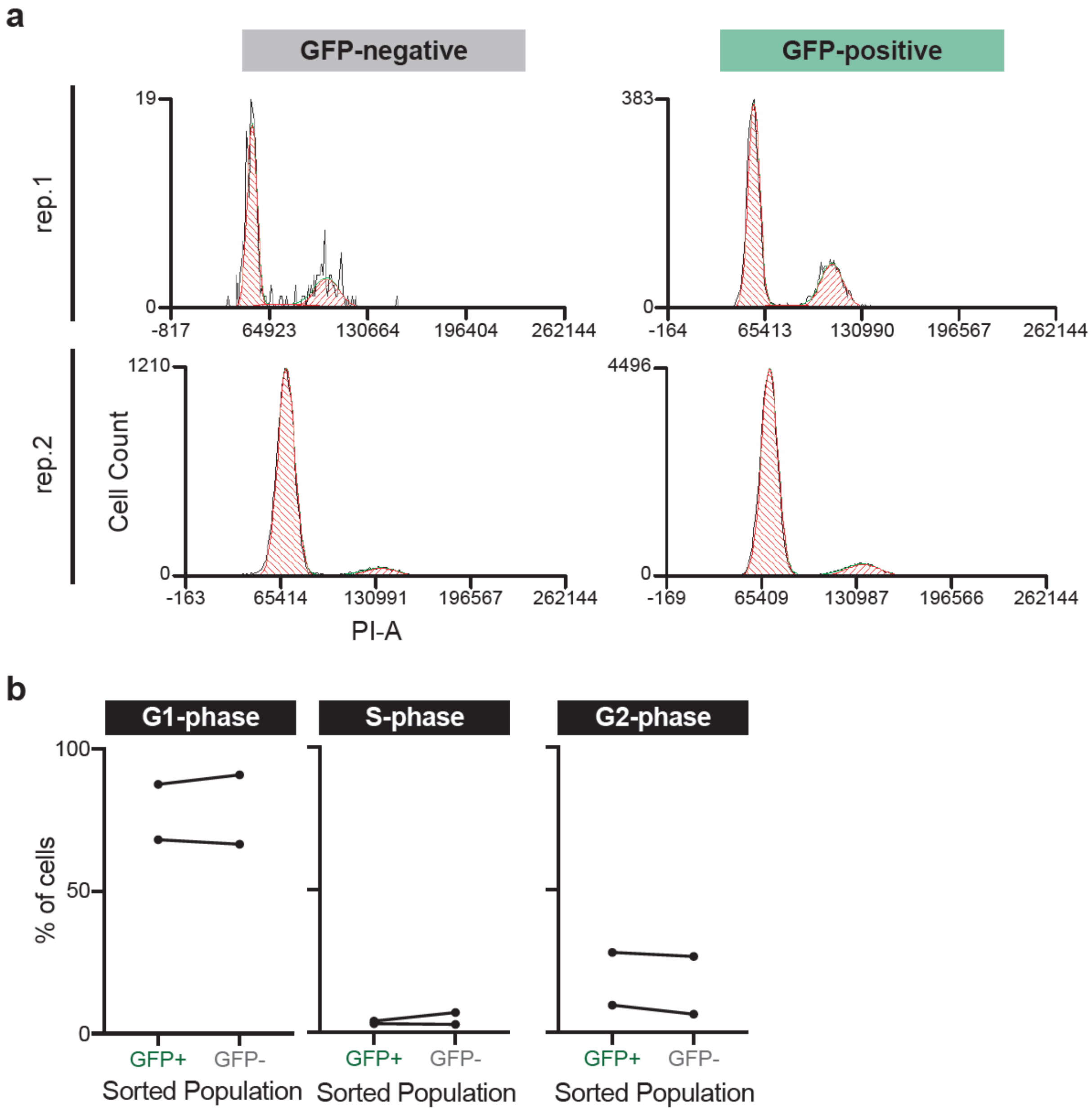


**Figure S9. READer activation is not cell cycle dependent.** NIH3T3 READer cells were plated and, after 36 hours, switched to GF-free media for 12 hours. Cells were then subjected to a 1 h pulse of 10% serum, incubated for 4 hours, and fixed and stained with propidium iodide (PI) for DNA content analysis of the GFP-high and GFP-low READer subpopulations. (**a**) Raw PI distributions, indicating cells' DNA content, are shown for the GFP-high and GFP-low subpopulations. Fits of the G1, S and G2 distributions are shown for two biological replicates. (**b**) Quantification of the percentage of cells in the different cell-cycle phases for both replicates, demonstrating negligible differences between GFP-high and GFP-low subpopulations.

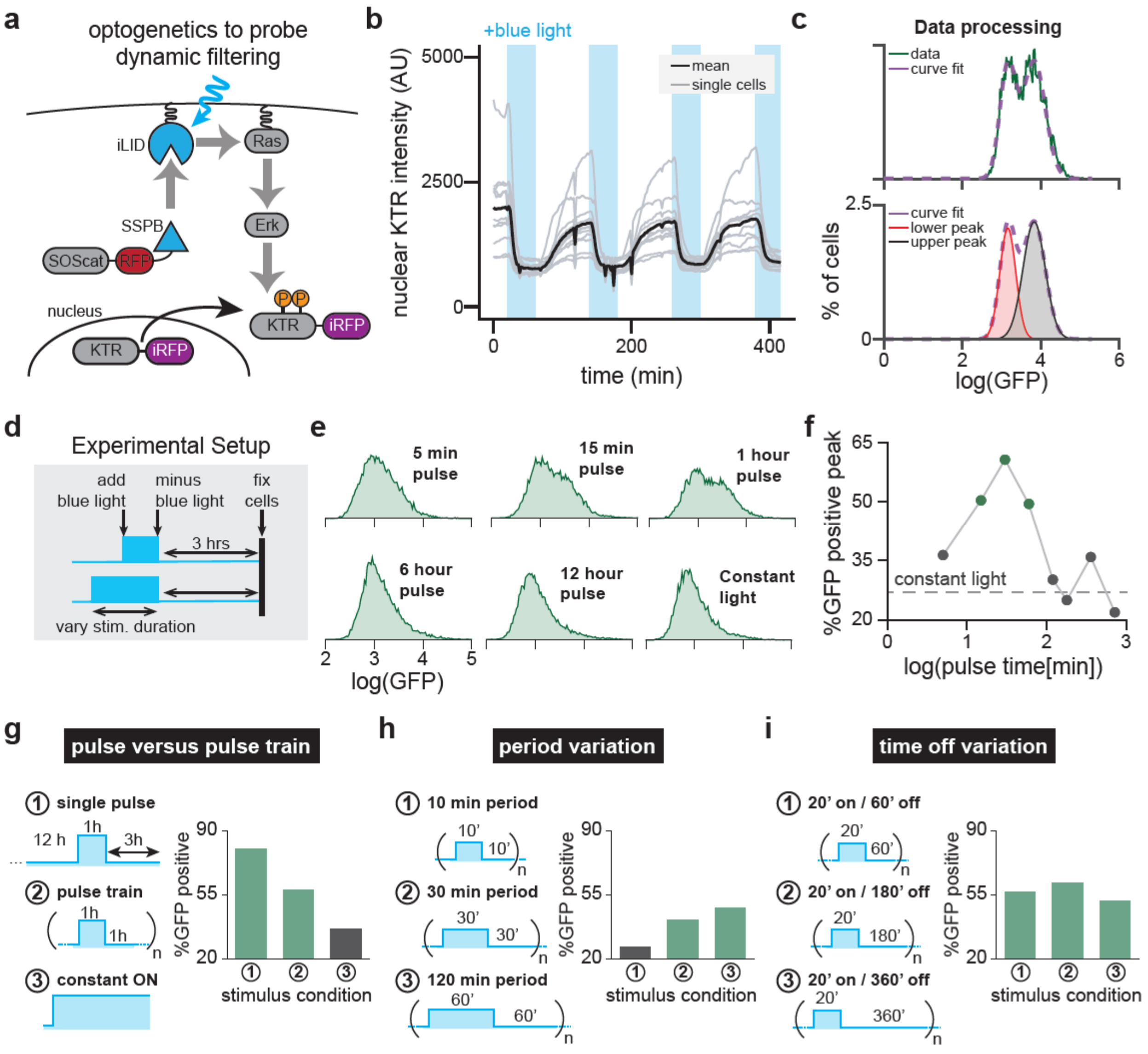


**Figure S10. OptoSOS enables characterization of READer dynamic filtering.** (**a**) Diagram of the OptoSOS system. Blue light stimulation heterodimerizes membrane-localized iLID-CAAX and TagRFP-SSPB-SOS$^{cat}$. Upon recruitment to the membrane, SOS$^{cat}$ activates Ras/Erk signaling. Upon phosphorylation by Erk, the ErkKTR-irFP biosensor is exported from the nucleus. (**b**) Single-cell traces showing nuclear KTR-iRFP during cycles of optogenetic stimulation. Blue bars shown when the DMD was turn on, providing cells with blue light. Shown are single cells (gray lines) and the mean of all cells (black line). (**c**) Data processing pipeline. The GFP histogram is fit to a sum of two Gaussians (top) and then the area under the curve of each peak is to approximate the fraction of GFP-high cells (bottom). (**d-e**) Mapping how optogenetic pulse stimulation affects READer circuit output. Light inputs of varying duration were applied to cells, which were fixed 3 h after the end of the pulse (schematic in **d**) and analyzed by flow cytometry for GFP induction (data in **e**). (**f**) Quantification of flow cytometry data in **e** reveals that 15-60 min pulses induce GFP production, while all other points are near the constant-on control (grey dotted line). (**g**) READer responses to single pulses versus pulse trains. Cells were given (1) a single 1 h light pulse, followed by 3 h in darkness before fixation, (2) a pulse train of alternating 1 h on/1 h off periods for 16 h or (3) constant light for 16 h. Quantification revealed that READer responds to both pulse trains and single pulses. (**h**) READer responses at various oscillation periods. Cells were given pulse trains at various periods $T$, each at 50% duty cycle, for 12 h: (1) $T = 20$ min, (2) $T = 60$ min or (3) $T = 120$ min. Case 1 resulted in low GFP, likely because the fast on/off cycles were interpreted by the cell as a constant stimulus, but Cases 2-3 led to GFP induction. (**i**) READer responses to various times between pulses. Repeated 20 min pulses were delivered every 60 min (case 1), 180 min (case 2) or 360 min (case 3) for a total of 12 h, with each leading to comparable GFP responses.

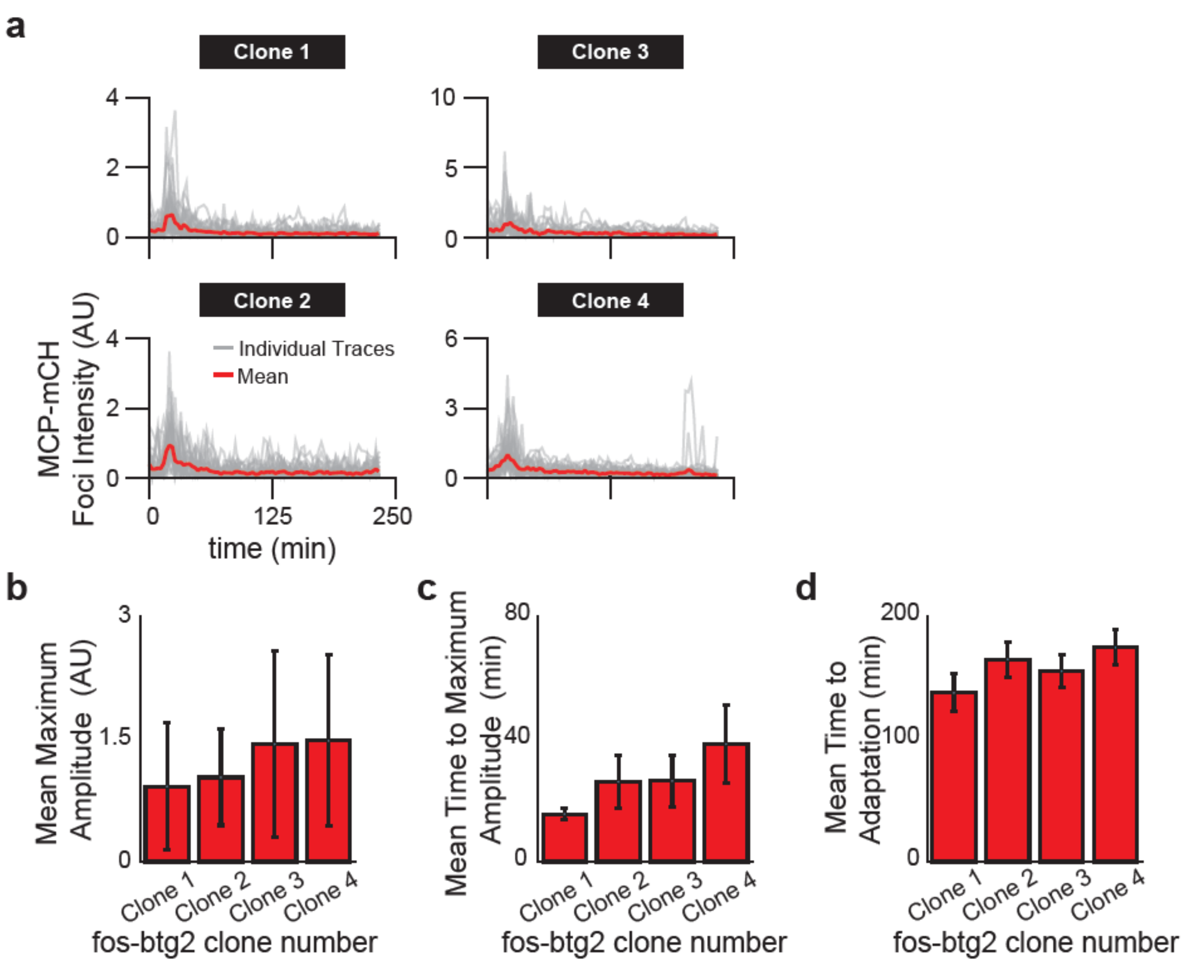


**Figure S11. Transcriptional adaptation from the *FOS* promoter in response to constant-on Erk stimuli.** We generated NIH3T3 cell lines expressing MCP-mCherry and where the PiggyBAC transposase was used to integrate a synthetic Erk target gene FOSp-YFP-24xMS2-3'*btg2*. Briefly, this construct contains the FOS promoter driving YFP, followed by 24 MS2 stem-loops and the *BTG2* 3' UTR, a long UTR for ensuring high MS2/MCP signal intensity. We then quantified transcriptional foci after treatment with 10% serum for cells in 4 independent clones derived from the parental cell line. (**a**) Quantification of individual foci (grey) and the mean (red) are shown for clone 1 (20 foci, 8 cells), clone 2 (12 foci, 4 cells), clone 3 (13 foci, 6 cells) and clone 4 (21 foci, 7 cells). (**b-d**) Analysis of transcription dynamics for foci shown in **a**. Parameters quantified include the mean maximum amplitude of foci (± S.D.) (in **b**), the delay time to maximum amplitude (in **c**) and the mean time to adaptation defined as the time when signal decays back down to 25% of maximum amplitude (in **d**). All error bars show ± S.E.M., except for **b** which shows ± S.D.

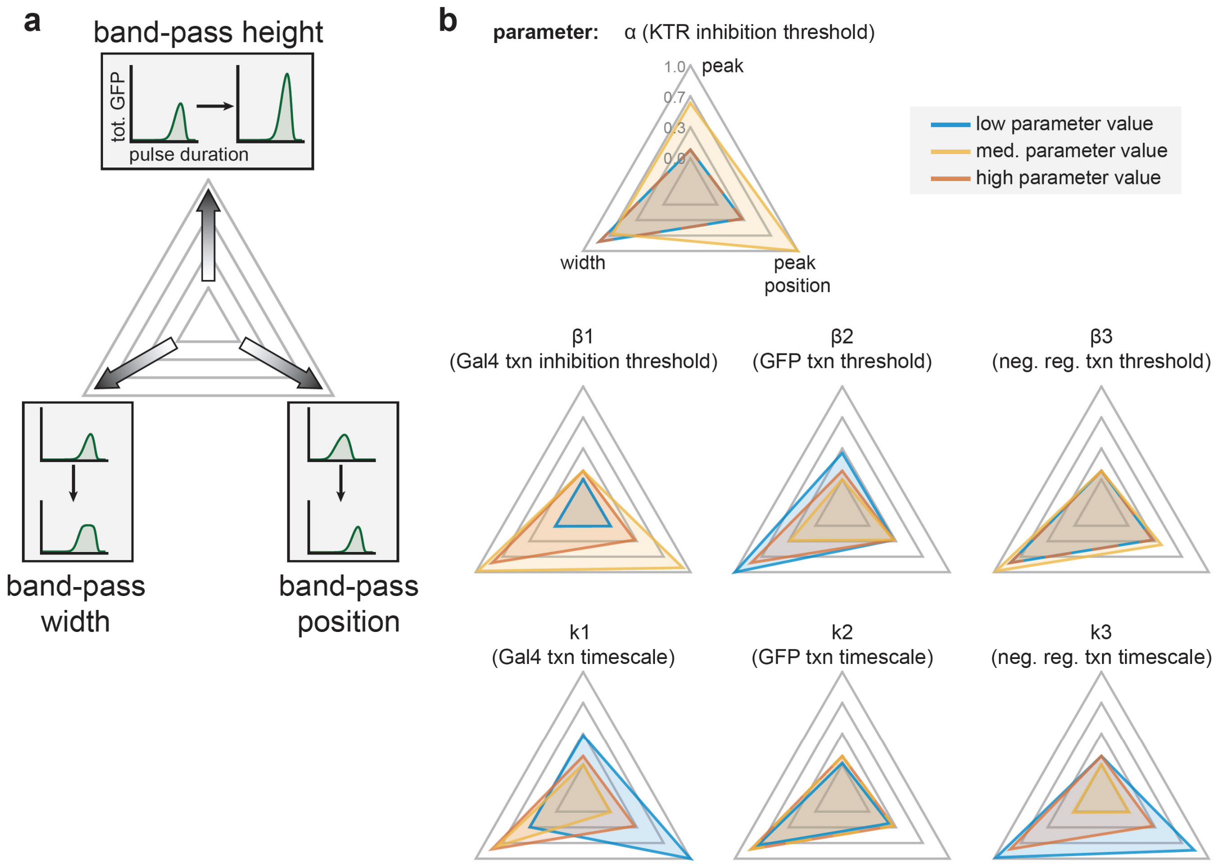


**Figure S12. Band-pass sensitivity analysis.** (**a**) For the mathematical model described in **Figure 2H**, each parameter was varied from 100-fold up and down the baseline value. The resulting band-pass curves were then analyzed for three features: band-pass height (maximum GFP response across all pulse durations), band-pass width (the span of durations achieving half-maximal GFP response) and band-pass position (the pulse duration that results in maximal GFP). (**b**) Radar charts showing the results of the parameter scan for the relative changes in all three band-pass features. The lowest, baseline and maximum values of the parameter are shown in blue, yellow and red, respectively. The further away from the center, the higher the value a particular simulation has for that feature. For example, parameters k1, k3, and β2 exhibit the highest shifts toward the lower-left corner, indicating their utility for tuning the READer circuit's selectivity (which we define as the width of the band-pass filter).

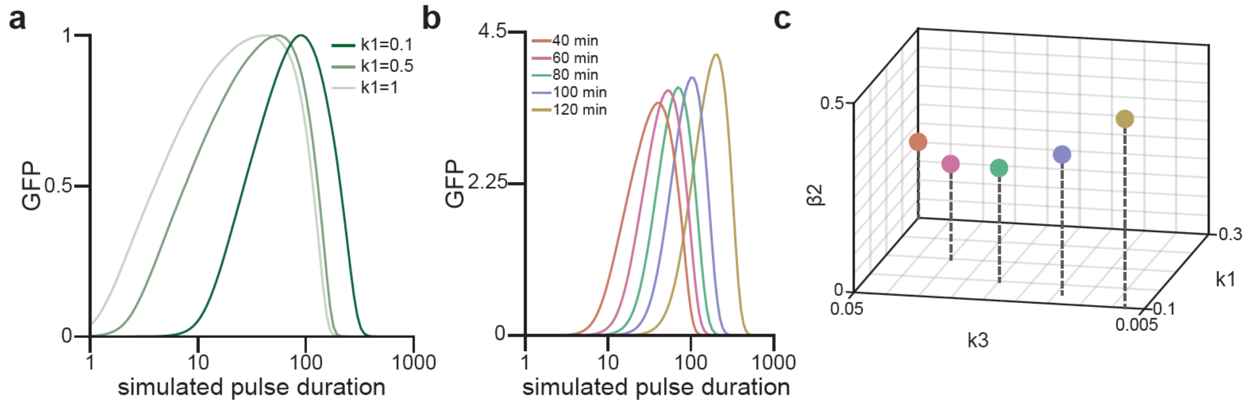


**Figure S13. The incoherent FFL with negative feedback implements a tunable band-pass filter.** (**a**) Simulated band pass curves (GFP output for pulses of different duration) produced for different values of the $x_1$ node's stability (parameter k1). The model predicts that destabilizing $x_1$ leads to better rejection of long pulses, which we later test experimentally (**Figure S14**). (**b**) Simultaneously tuning all three parameters (k1, k3, and b2) enables more complete control over band-pass filtering. Here we show simulated band pass curves at 5 distinct optimal pulse durations, but with similar band pass widths and amplitudes, achieved by tuning all three parameters. (**c**) Plotted are the parameters used to construct the band-pass curves in **a**. If one follows this curve in parameter space by tuning the timescale of negative feedback strength, $x_1$'s production, they can develop a filter that transmits ("passes") pulses of defined length.

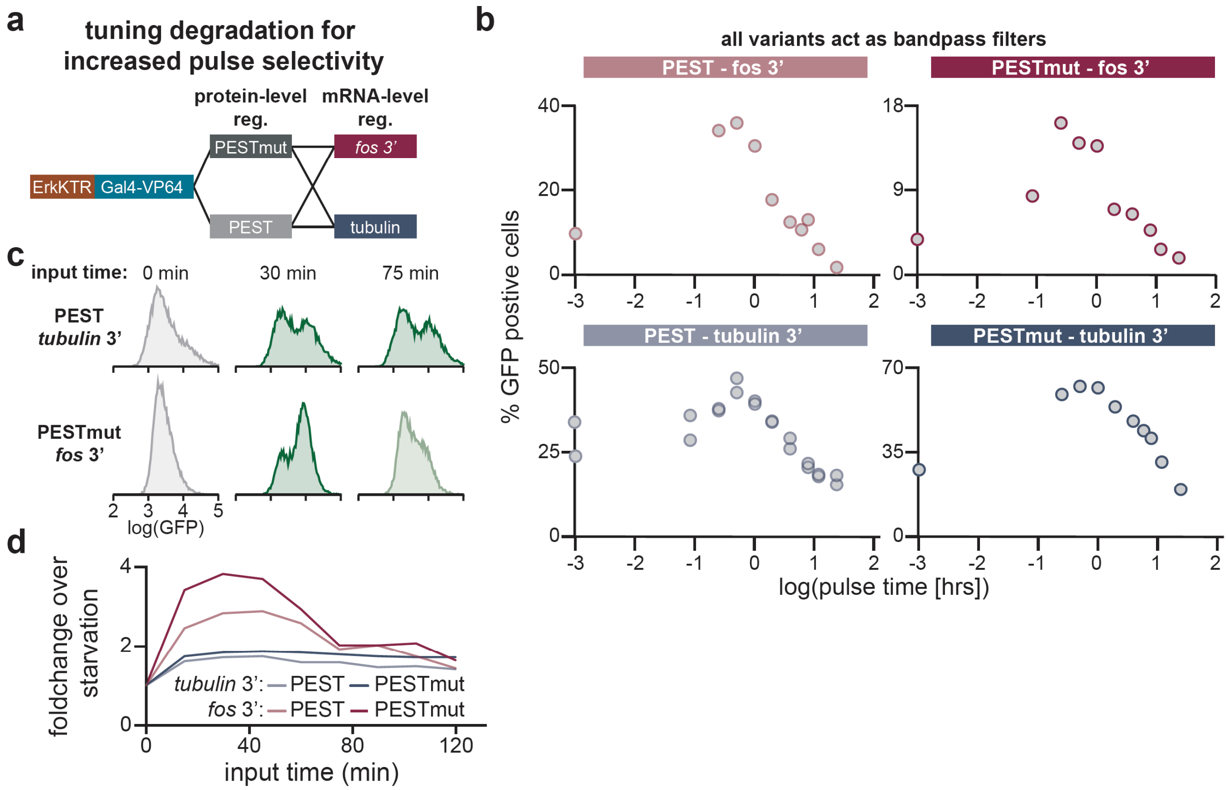


**Figure S14. Modulating KGV degradation modulates pulse filtering.** (**a**) To change the stability of the KTR-Gal4-VP64 transcription factor we replaced the PEST sequence with a mutated one with a decreased half-life (PESTmut), changed the *tubulin* 3' UTR to the *fos* 3' UTR which is more unstable, or both. All variants were integrated into the same reporter cell line that contained the UAS-GFP construct as well as OptoSOS. (**b**) Quantification of GFP flow cytometry results from all variants in response to different pulse lengths and a three hour off period. (**c**) Representative GFP histograms for the PEST-*tubulin* 3' UTR and PESTmut-*fos* 3' UTR variants in response to constant dark (starvation), 30 minutes of blue light or 75 minutes of blue light. PEST-*tubulin* 3' UTR variant responds to both pulses while the PESTmut-*fos* 3' UTR variant only responds to the shorter pulse. (**d**) Expanded analysis of **c** where all variants were given pulse durations from 0 minutes to 2 hours on a 15 minute interval. Either variant with the destabilized *fos* 3' UTR results in more selective pulse filtering.

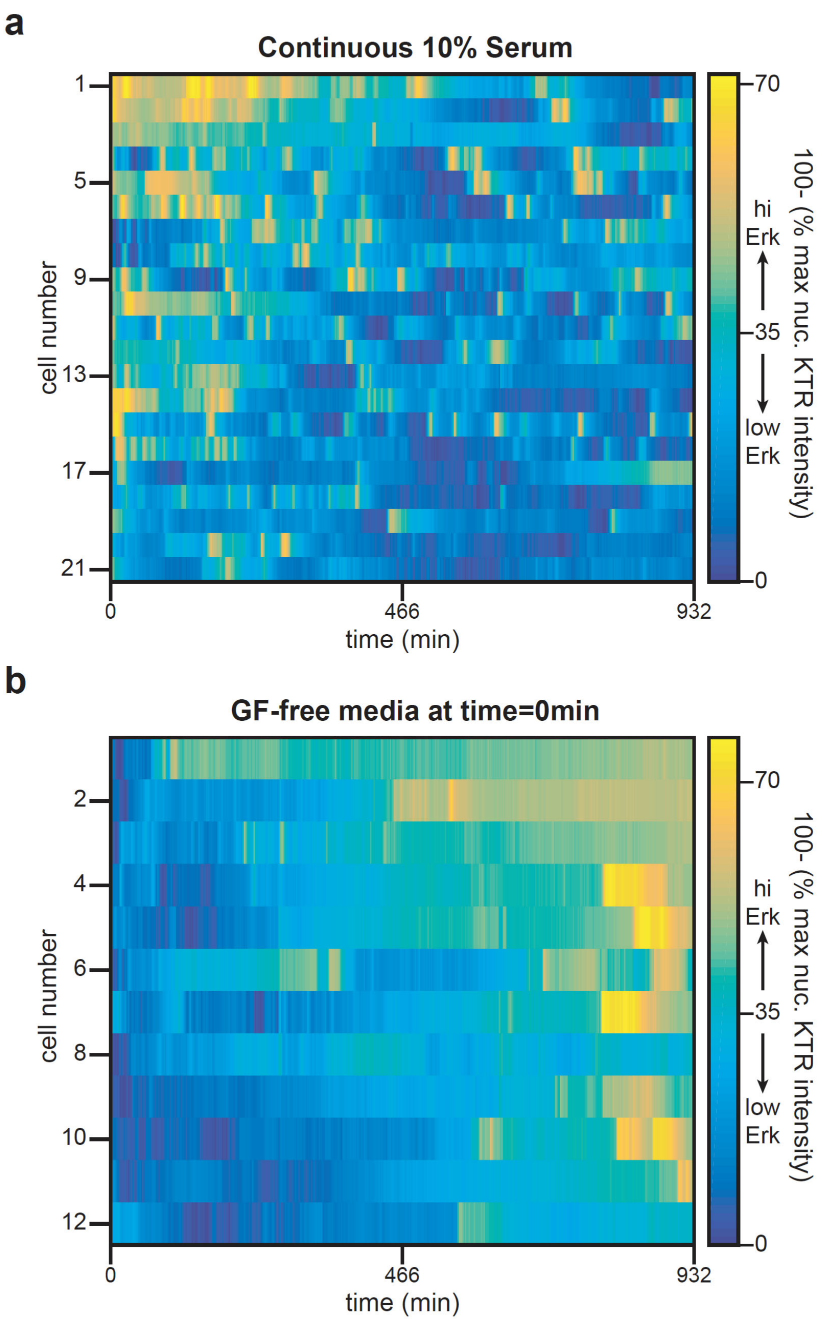


**Figure S15. Erk dynamics in NIH 3T3s.** NIH 3T3s expressing Erk-KTR-iRFP were plated in 96-well plate wells and either left in growth media for the entirety of imaging (**a**) or switched into growth-factor free media at the beginning of imaging (**b**). Nuclear iRFP fluorescence was quantified for 21 and 12 cells for the two cases, respectively. Heatmaps show these single cell traces for (100 – the nuclear intensity normalized to each cell's maximum nuclear fluorescence).

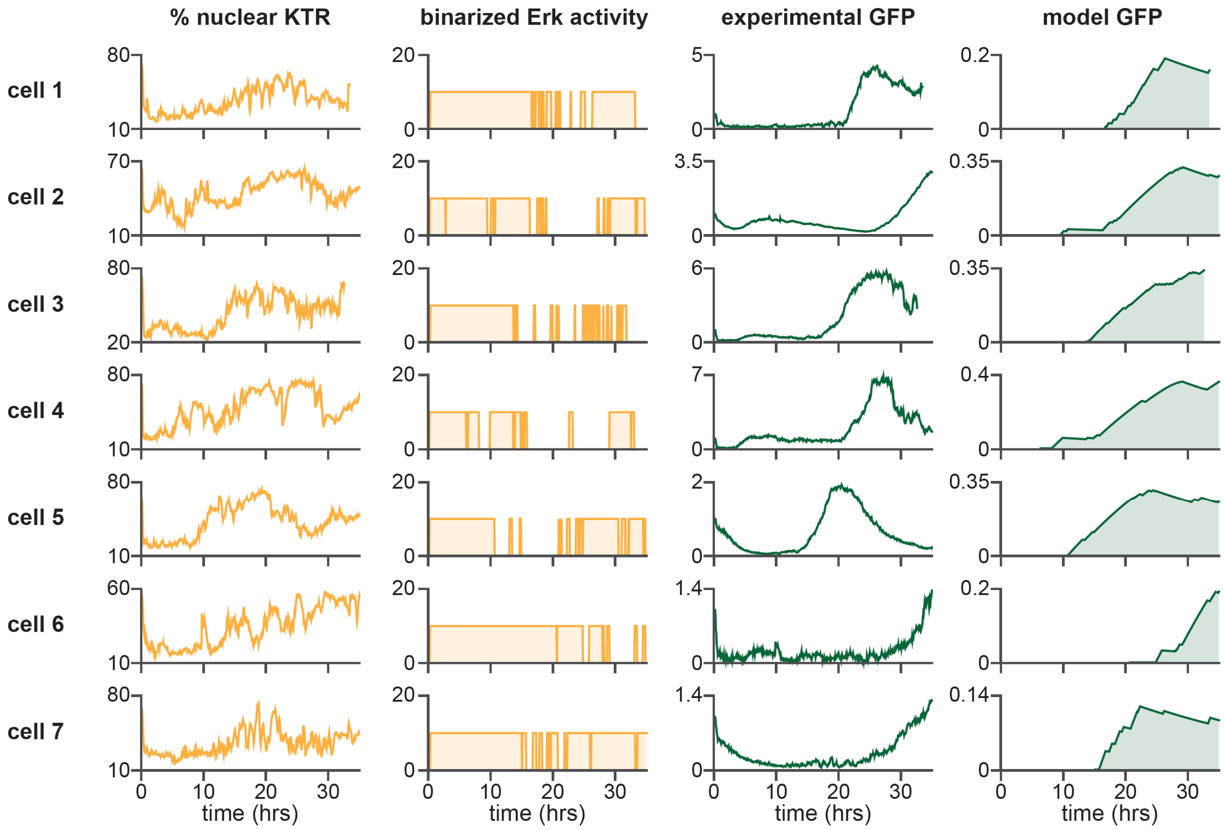


**Figure S16. Additional cells for capturing endogenous pulses in NIH 3T3s.** Quantification from representative READer cells expressing KTR-mScarlet stimulated with serum at time 0 and imaged for 34 hours. From left to right, columns indicate quantification of (1) % nuclear KTR, (2) "binarized" KTR traces based on thresholding the % nuclear KTR trajectories, (3) fold-change in GFP intensity and (4) simulated GFP response when the binarized KTR-mScarlet trace is used as a model input. Rows represent analyses for 7 different cells. Cell 1 is the same as shown in **Figure 3C**.

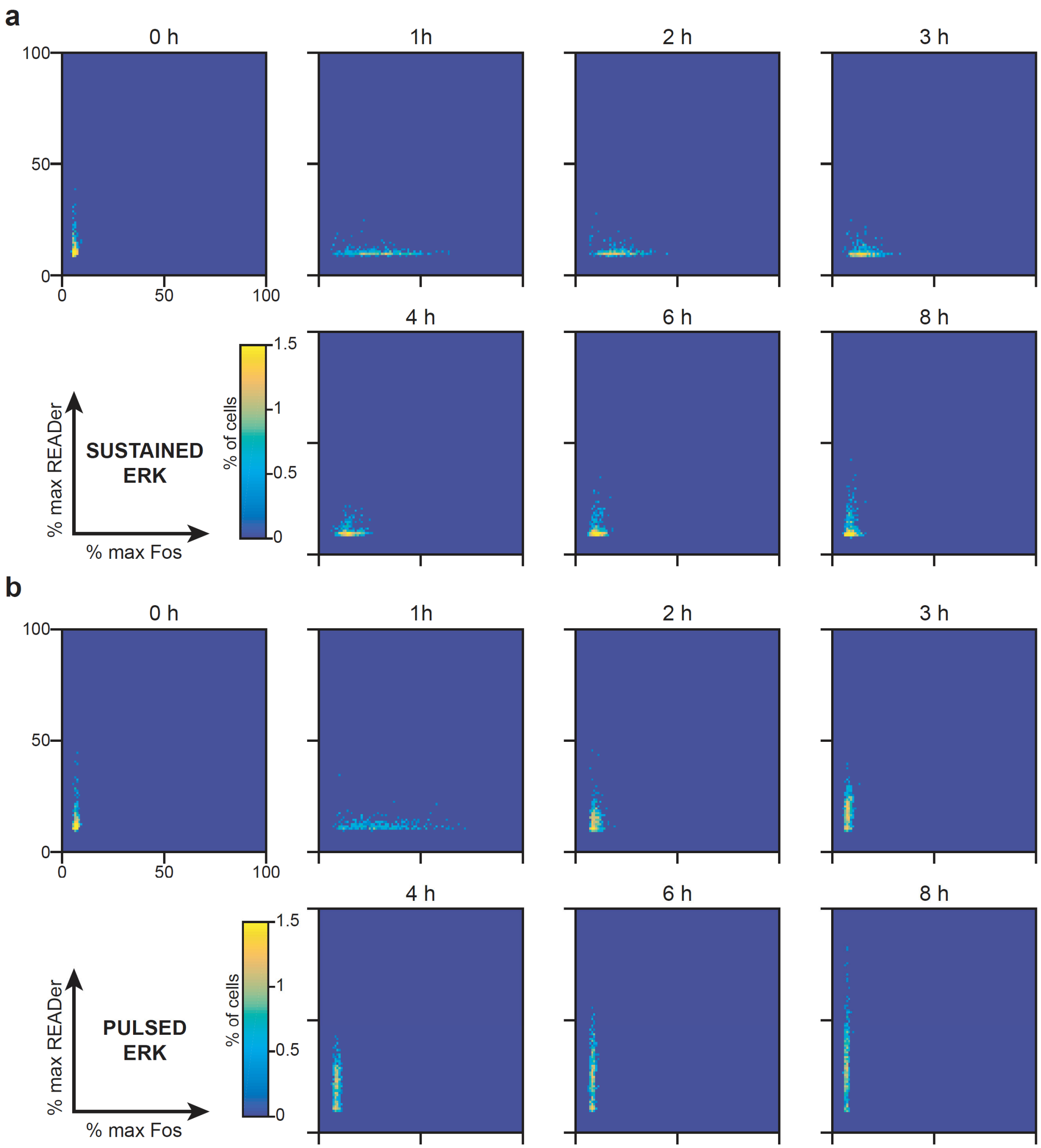


**Figure S17. Density plot analysis of Fos-READer immunofluorescence experiment.** Quantification of immunofluorescence intensity from fixed cells for Fos and READer for (**a**) sustained serum stimulation or (**b**) a 20-minute pulse of serum. Each cell's intensity is normalized to the maximum Fos or READer GFP intensity for the experiment and then plotted on the Fos-READer plane where Fos is the horizontal axis while READer is the vertical axis. Each plot represents a different timepoint after the beginning of stimulation. In both stimulation conditions, Fos activation occurs at the 1-hour timepoint. Fos stays high in the sustained input case but goes down rapidly in the pulsed case. Only in the pulsed case is there any activation of READer GFP.

#### Supplementary Movie Legends

**Movie S1. Time-lapse imaging of NIH3T3 READer cells in response to different dynamic stimuli.** TagRFP (white) is uniformly expressed in READer cells and can be used to identify cells. GFP (green) reports on READer system activation. Cells were imaged with a 60X oil objective every 6 min for 5 h. Panels show three stimuli: constant GF-free media (constant OFF; left), constant 10% serum (constant ON; middle), and a 10% serum pulse for 1 hour followed by GF-free media (pulse; right).

**Movie S2: Long-term imaging of NIH3T3 cells expressing ErkKTR-iRFP reveals spontaneous pulses.** Cells were imaged using a 20X air objective every 3 min for 23 h in either growth media containing 10% serum (left) or were switched to GF-free media immediately prior to imaging (right).

**Movie S3: Time-lapse imaging of representative NIH3T3 READer cell expressing ErkKTR-mScarlet.** Cells were imaged using a 20X air objective every 3.5 min for 48 h. After having been placed in GF-free media overnight, cells were stimulated with constant 10% serum at the beginning of imaging. Images show ErkKTR-mScarlet (left) and GFP (right), which report on Erk dynamics and READer system activation, respectively. Quantification of the images are shown below each, with the fraction of nuclear KTR shown on the left, and the GFP fluorescence intensity shown on the right.
